# Genome reduction increases endosymbiont sensitivity and promotes mutualism evolution

**DOI:** 10.64898/2026.10.08.756896

**Authors:** Yuki Kanai, Chikara Furusawa

## Abstract

Endosymbiosis is a major source of biological innovation, enabling hosts to explore new ecological niches. While genome reduction in endosymbionts is well documented, its impact on the evolutionary transition from parasitism to mutualism remains unclear. Regulatory genes are lost disproportionately as bacterial genomes shrink, as we confirm in endosymbiont-containing lineages. We therefore model metabolic competition between hosts and endosymbionts, representing endosymbiont genome size by the number of independently regulated sets of reactions. In our simulations, hosts that grow poorly alone adapt best when harboring moderately genome-reduced endosymbionts rather than large-genome ones. This advantage goes beyond the lower resource use of smaller endosymbionts. By decomposing endosymbiont metabolic responses to changes in host-provided metabolites into “adaptability” and “sensitivity,” we demonstrate that moderate genome reduction increases the sensitivity of endosymbionts to host metabolic conditions, rendering them more exploitable through host evolution. This suggests that genome reduction plays a more constructive role in mutualism evolution than previously appreciated, facilitating host-level selection for exploitable endosymbionts rather than being merely a passive consequence of endosymbiosis. Our systems-level approach, focusing on regulatory capacity rather than specific gene functions, provides a general framework for understanding how genome reduction affects the evolutionary trajectory from parasitic to mutualistic endosymbionts, and may help explain the frequent replacement of highly reduced endosymbionts in nature.

## Introduction

Many eukaryotes, including plants and animals, harbor endosymbiotic microorganisms that are vertically transmitted from parent to offspring. These partnerships have been transformative: organelles such as mitochondria and plastids are ancient bacterial endosymbionts that became integral to eukaryotic life, and more recent endosymbiotic acquisitions are found in diverse eukaryotic lineages, including protists, fungi and metazoans (Keeling and McCutcheon, 2017). Such endosymbiotic acquisitions have enabled hosts to access new ecological niches, driving the diversity and success of eukaryotes (Nowack and Melkonian, 2010; Sudakaran et al., 2017; Cornwallis et al., 2023).

Despite these benefits, the evolution of mutualistic endosymbiosis presents a fundamental challenge. Unlike horizontally transferred genes, which hosts can directly regulate, endosymbionts are autonomous organisms that have evolved to maximize their own fitness. Indeed, comparative studies indicate that mutualistic endosymbionts often evolved from parasitic ancestors (Sachs et al., 2014). This poses the following problem: how do hosts transform initially selfish and often parasitic endosymbionts into beneficial partners? Theory suggests that vertical transmission facilitates mutualism evolution by aligning host and endosymbiont evolutionary interests (Frank, 1996), and that hosts can stabilize mutualism by choosing, rewarding or punishing partners (Bull and Rice, 1991; Kiers et al., 2003; Sachs et al., 2004). These frameworks explain how mutualistic partners are favored and maintained, but largely assume that such partners are available to be selected. How mutualistic variants arise from initially selfish endosymbionts remains elusive (Drew et al., 2021), because the early stages of this transition are rarely observed in nature.

We propose that genome reduction—the most conspicuous consequence of the evolution of endosymbiosis—can facilitate the evolution of mutualism. Endosymbionts typically undergo extreme genome reduction, shrinking from the megabase-sized genomes of free-living relatives (Mira et al., 2001), in some lineages to just hundreds of kilobases (McCutcheon and Moran, 2012). This reduction is commonly viewed as passive evolutionary decay due to relaxed selection: stable environments rendering genes superfluous, bottlenecks weakening purifying selection to retain genes, and isolation within the host body preventing horizontal gene acquisition (O’Fallon, 2008; McCutcheon and Moran, 2012).

If genes were lost randomly during genome reduction regardless of function, the relative abundance of each functional category would be expected to remain approximately constant as genomes shrink. However, this is not the case. Across bacterial genomes, genes in different functional categories scale differently with genome size: genes involved in information processing, such as translation, tend to be lost more slowly and scale sublinearly; metabolic genes scale approximately linearly; and regulatory genes scale superlinearly, in some datasets approaching a quadratic relationship (van Nimwegen, 2003; Sela et al., 2019). This relationship is also observed in Enterobacterales, an order that contains known endosymbionts at different stages of genome reduction (Fig. 1A). The ratio of regulatory to metabolic genes decreases with decreasing genome size both across Enterobacterales (Fig. 1A) and within several endosymbiont-containing lineages (Fig. 1B), indicating disproportionate, and potentially preferential, loss of regulatory genes during genome reduction.

**Figure 1.**
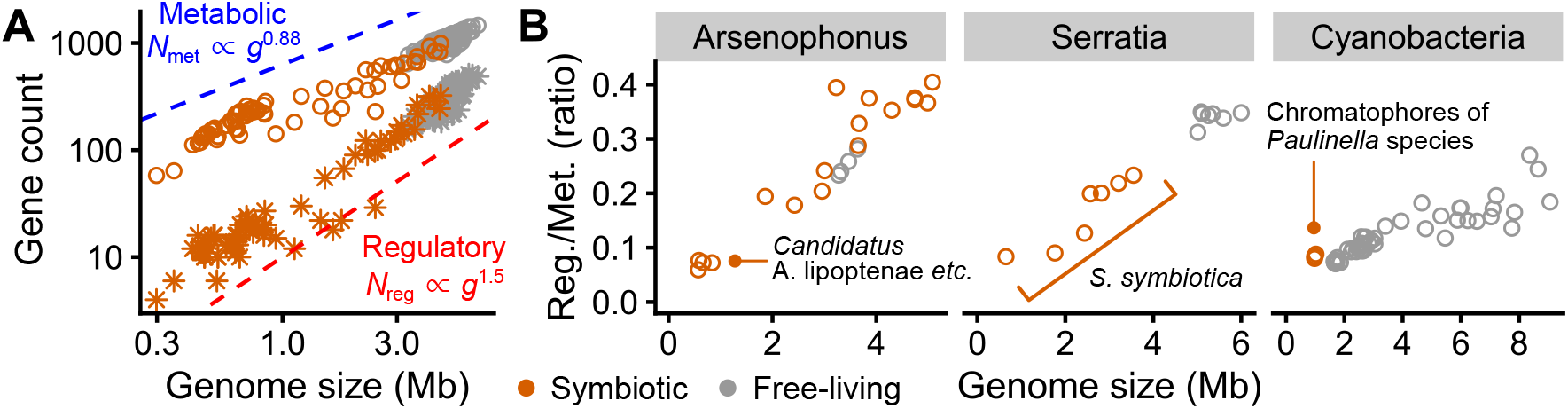
The rapid loss of regulatory genes. **(A)** Global scaling relationships between genome size (*g*) and the numbers of regulatory and metabolic genes in genomes (*N*_reg_, *N*_met_) from the order Enterobacterales, which includes organisms at various stages of endosymbiotic genome reduction (genomes chosen as in (Forni et al., 2026)). Regulatory genes decline more rapidly with decreasing genome size than metabolic genes, as indicated by the steeper regression slope for regulatory genes (red) than for metabolic genes (blue). **(B)** The ratio of regulatory to metabolic genes also decreases with decreasing genome size within closely related endosymbiont lineages. Facets, from left to right, show *Arsenophonus* (Siozios et al., 2024), *Serratia* (Manzano-Marín and Latorre, 2016; Forni et al., 2026) and cyanobacteria closely related to chromatophores of *Paulinella*, including members of the genera *Synechococcus* and *Prochlorococcus* (Larsson et al., 2011; Lhee et al., 2019). Methods for data collection and analysis are described in *SI Appendix*, section S1. Raw data are available in Dataset S1.

Consistent with this, regulatory genes have been reported to be preferentially lost (McCutcheon and Moran, 2012; Galán-Vásquez et al., 2016; Nechitaylo et al., 2021), and such losses can directly promote mutualistic interactions (Koga et al., 2022; Gabr et al., 2022). Remarkably, the genome of *Buchnera aphidicola*, an aphid endosymbiont, is essentially a subset of the genome of the free-living model organism *Escherichia coli* (Shigenobu et al., 2000), suggesting that mutualism can emerge through changes in gene regulation rather than through acquisitions of novel “symbiosis” genes (Moran, 2003). Together, these findings suggest that the preferential loss of regulatory capacity is not merely evolutionary decay, but may create conditions that facilitate host-level selection for mutualism.

Here, we test the hypothesis that the loss of regulatory genes can make endosymbionts more amenable to host control. While previous studies have primarily focused on specific genes lost during genome reduction in particular endosymbionts, we investigate the general impact of genome reduction, which we model as a decrease in the endosymbiont’s capacity to regulate its metabolism. To do this, we adopt a computational model of nutritional symbiosis between a host and an endosymbiont, each with a randomly generated metabolic network. Simulating how hosts adapt to challenging environments by acquiring endosymbionts with varying genome sizes reveals that genome reduction facilitates the evolution of mutualism beyond simply reducing parasitism. Our key finding is that the metabolism of genome-reduced endosymbionts shows increased “sensitivity” to changes in the host’s internal environment. This increased sensitivity creates greater phenotypic variation among endosymbionts in response to different host genotypes, leading to more diverse metabolic exchanges and variable fitness outcomes among hosts. This enhanced variation gives host-level selection more beneficial variants to act on, facilitating the evolutionary transition toward mutualism.

## Results

### Modeling the metabolic competition between hosts and endosymbionts

We model the evolution of nutritional mutualism after host organisms acquire new endosymbionts with varying genome size, represented in the model by the complexity of their gene regulation (Fig. 2A). We consider a situation in which hosts acquire endosymbionts in environments where they grow poorly on their own, for example because a key metabolite is scarce but can be produced by the endosymbiont. At the time of acquisition, however, the endosymbiont is not assumed to be beneficial to the host. Instead, because the endosymbiont autonomously maximizes its own growth, it may initially behave as a parasite by consuming metabolites that would otherwise support host growth. We therefore model the early host–endosymbiont relationship as metabolic competition for shared metabolites within the host body and assume that each organism maximizes its growth by regulating its metabolism.

**Figure 2.**
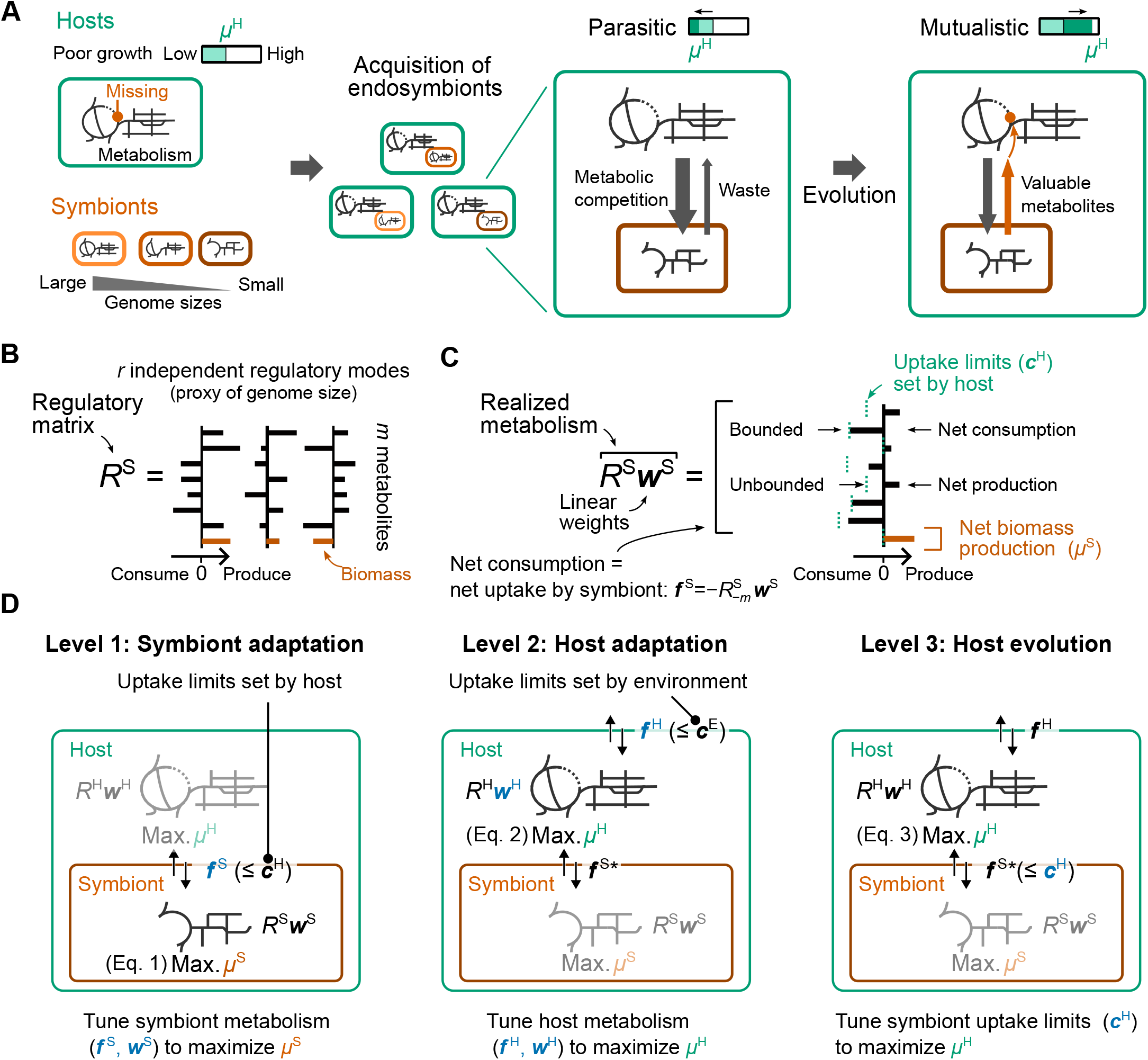
Overview of the model. **(A)** To study the impact of endosymbiont genome size on the evolution of mutualism, we randomly assigned each host an endosymbiont of varying genome size (left), and the host evolved to improve its growth in the presence of the endosymbiont (right). We modeled a nutritional symbiosis where hosts and endosymbionts compete for the same *m* metabolites, and hosts evolve to control endosymbionts by tuning the metabolites available to them, so that endosymbionts end up producing metabolites that are beneficial to hosts. **(B)** Both endosymbionts and hosts optimize their metabolism by tuning the weights of their regulatory modes, which are combinations of metabolic reactions regulated by a shared regulatory gene. Each organism has a regulatory matrix (*R*^S^ for endosymbionts and *R*^H^ for hosts) that represents the regulatory modes of the organism’s metabolism. As a proxy for genome size, we randomly generated *R*^S^ with varying numbers of columns or regulatory modes *r*, which reflects the complexity of endosymbiont gene regulation. Among the *m* metabolites, the *m*-th metabolite is considered the sole biomass precursor. **(C)** To maximize their growth (*µ*^S^), endosymbionts adjust the metabolic conversions they perform via nonnegative weights assigned to each regulatory mode (***w***^S^). Following the conventions of constraint-based modeling, any net consumption of non-biomass metabolites must be balanced by their uptake from the host, which cannot exceed the uptake limits ***c***^H^. Thus, in the figure, bars corresponding to non-biomass metabolites in the left direction indicate net consumption or uptake, and bars in the right direction indicate net production or excretion by the endosymbiont. Following the conventions of linear programming, we refer to metabolites that are consumed up to their host-provided limit as “bounded” metabolites, and all others as “unbounded” metabolites. Hosts adjust their metabolism in a similar manner, with their uptake rates from the environment bounded by ***c***^E^ and with the metabolites consumed or excreted by their endosymbionts (***f*** ^S∗^) entering their mass balance. **(D)** The model consists of three levels of optimization as in Eqs. 1–3, from fastest to slowest: symbiont adaptation, host adaptation and host evolution. Each level takes the optimum of the level below as given; gray elements are held fixed within a level. See *SI Appendix*, section S3 for the full formulation, and *SI Appendix*, Table S1 for the list of model parameters. For brevity, endosymbionts are labeled “symbionts” in the figures.

To focus on the general consequence of genome reduction rather than on specific gene contents or network architectures, we developed a toy model with randomly generated regulatory matrices that define the metabolic conversions available to hosts and endosymbionts and how these conversions are regulated.

Host–endosymbiont pairs interact through *m* metabolites, denoted as *M*_1_, …, *M*_*m*_. Unless otherwise stated, we set *m* = 10; however, the exact value of *m* does not qualitatively affect the main results, as discussed below. For simplicity, we designated *M*_*m*_ as the only biomass metabolite. We assume that biomass production is the sole determinant of fitness for both hosts and endosymbionts. Metabolic conversions were limited to one-to-one conversions between metabolites, such as *M*_*i*_ →*M*_*j*_. For each organism, possible one-to-one conversions were generated from the _*m*_C_2_ = 45 metabolite pairs, with directions randomized across generated networks.

To express the regulation of metabolism, we assume that the metabolism of each organism is controlled by tuning the weights of regulatory modes, which are combinations of metabolic reactions regulated by a shared regulatory gene. Examples include genes in the same operon or genes that are coregulated by the same transcription factor. The set of regulatory modes is represented by a regulatory matrix *R*^X^, where X is either H (host) or S (endosymbiont) (Fig. 2B). Each column of the matrix denotes a “regulatory mode.” If reaction A converts metabolite *M*_1_ to *M*_2_ (A: *M*_1_ →*M*_2_), this reaction is expressed as the vector ***v***_*A*_ = (−1, 1, 0, …, 0)^⊤^, where the first element expresses the consumption of metabolite *M*_1_ and the second element expresses the production of metabolite *M*_2_. If the *j*-th regulatory mode upregulates reaction A as well as reaction B, which converts *M*_2_ into the biomass metabolite *M*_*m*_ (B: *M*_2_ →*M*_*m*_), the regulatory mode is expressed as the vector ***q***_*j*_ = *κ*_*jA*_***v***_*A*_ +*κ*_*jB*_***v***_*B*_, where *κ*_*jA*_ and *κ*_*jB*_ are positive weights representing the relative contributions of reactions A and B to the regulatory mode.

Organisms regulate their metabolism by changing the nonnegative weights, or expression levels, of *r* different regulatory modes, as in *w*_1_***q***_1_ + *w*_2_***q***_2_ +· · · + *w*_*r*_ ***q***_*r*_. Equivalently, if the regulatory matrix is written as *R* = [***q***_1_, ***q***_2_, …, ***q***_*r*_], the realized metabolic conversion is given by *R****w***, where ***w*** = (*w*_1_, *w*_2_, …, *w*_*r*_)^⊤^ (Fig. 2C). For example, consider an endosymbiont with one regulatory mode ***q***_1_, defined as ***q***_1_ = ***v***_*A*_ +***v***_*B*_. If the host allows the endosymbiont to take up *M*_1_ at a rate of at most one unit, this endosymbiont can convert it into biomass at the same rate by upregulating reactions A and B, which is represented by assigning the mode weights ***w*** =(1)^⊤^. If the host instead allows uptake of *M*_3_ at a rate of at most one unit, the endosymbiont cannot convert it into biomass, because its regulatory mode does not include any reaction that connects *M*_3_ to biomass. However, if the endosymbiont has a second regulatory mode, ***q***_2_ = ***v***_*C*_, which upregulates reaction C (*M*_3_ →*M*_1_), the endosymbiont can convert *M*_3_ into biomass by expressing both regulatory modes, with weights, for instance, ***w*** = (1, 1)^⊤^. As illustrated by this example, having more regulatory modes allows the endosymbiont to adapt to a wider range of host-provided metabolites, increasing its autonomy.

Therefore, we used the number of regulatory modes *r* as a proxy for the genome size of the endosymbiont. A decrease in *r* in this model corresponds to decreases in the number of transcription factors that regulate metabolic genes (McCutcheon and Moran, 2012) and in the number of independent regulons due to increased operon density (Nuñez et al., 2013), both of which simplify endosymbiont regulation. Because each mode carries a set of reactions, a decrease in *r* reduces both the regulatory and the metabolic capacity of the endosymbiont, as occurs during endosymbiont genome reduction. For the host, we fixed the number of regulatory modes at *r*^H^ = 24, which is large enough relative to *m* = 10 that hosts can grow in many environments, but not so large that hosts never require the endosymbiont. We treat *r* as an exogenous parameter for most of this study and return to its own evolution below.

We randomly generated the regulatory matrices *R*^S^ and *R*^H^ (see *SI Appendix*), with the number of regulatory modes *r* in *R*^S^ ranging from 1 to 100. Each regulatory mode ***q***_*j*_ was assembled by drawing *k* = 4 reactions from the organism’s reaction pool and assigning them positive weights *k*_*jA*_, *k*_*jB*_, etc.

Given these regulatory matrices, we model the endosymbiotic relationship as three levels of optimization ordered by timescale (Fig. 2D). For simplicity, and to focus on the effect of genome size, we assume a complete separation of these timescales, so that each level takes the optimum of the faster levels as given and holds the slower levels fixed. The regulatory matrices, the set of available metabolic genes and the host– endosymbiont pairing are held constant throughout, on the assumption that they change on much longer timescales. This formulation captures how hosts, unable to directly control autonomous endosymbionts, can nonetheless evolve to exploit them through physiological change.

The first two levels express metabolic competition between the host and endosymbiont for resources within the host body. First, endosymbionts regulate their gene expression (***w***^S^) to maximize their own growth (*µ*^S^), converting metabolites supplied by the host, at rates up to ***c***^H^, into biomass. Second, hosts regulate their gene expression (***w***^H^) to maximize their own growth (µ^H^) while taking up metabolites from the environment at rates up to ***c***^E^ and exchanging metabolites with their endosymbionts (***f*** ^S^).

Third, hosts evolve to control their endosymbionts by altering the metabolites available to them. Hosts are known to regulate nutrient supply at the symbiotic interface, for example through amino acid transporters expressed in aphid bacteriocytes (Price et al., 2011, 2014), branched-chain amino acid supply to rhizobial bacteroids (Prell et al., 2009) and nitrogen availability to algal symbionts (Cui et al., 2019). Accordingly, we assume hosts can evolve to alter the maximum rates at which endosymbionts can take up each metabolite (***c***^H^) within their internal environment, such as the gut, cytoplasm, or specialized cells like bacteriocytes. To focus on the impact of genome reduction rather than on changes in ecological interactions, we assume that hosts have already established mechanisms to vertically transmit their endosymbionts to offspring, thereby sidestepping the need to model the evolution of acquisition mechanisms. This scenario most closely resembles genome reduction following symbiont replacement (Husnik and McCutcheon, 2016). We place hosts in environments where they reproduce poorly without endosymbionts and allow them to evolve by optimizing the metabolic environment they provide to their endosymbionts. This framework enables hosts to derive benefits from endosymbionts even when endosymbionts primarily maximize their own growth.

The mathematical formalization of the three levels is described below, and the full formulation of the trilevel optimization problem is provided in *SI Appendix*, Eq. S6. Following flux balance analysis, all quantities in the model are steady-state rates in arbitrary units: ***c***^H^ and ***c***^E^ are upper bounds on the rates at which endosymbionts and hosts, respectively, can take up each metabolite. We refer to them as uptake limits, and to ***c***^H^ as the host’s internal environment, which determines the host-provided metabolites.

1. **Symbiont adaptation:** Endosymbionts adapt to maximize their growth (*µ*^S^) given the maximum rates at which they can take up each metabolite from the host (***c***^H^) (Fig. 2D). To maximize biomass production (*µ*^S^), endosymbionts regulate their metabolism by controlling two factors: the activation levels or weights of their regulatory modes (***w***^S^) and the net uptake of metabolites from the host (***f*** ^S^). This optimization is constrained by the uptake limits set by the host (***f*** ^S^ ≤***c***^H^). Following previous constraint-based analyses of endosymbiont metabolism (Pál et al., 2006; Thomas et al., 2009; Ankrah et al., 2018), we formulate this optimization as a linear programming problem:

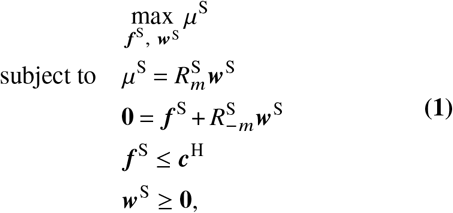

where the subscripts *m* and −*m* denote the *m*-th row and all other rows, respectively. The first equality expresses biomass production. The *m*-th row of the regulatory matrix *R*^S^ corresponds to the quantity of biomass produced per unit of the regulatory mode expressed (Fig. 2B). The right-hand side expresses the weighted sum of the regulatory modes, leading to net biomass production *µ*^S^ (Fig. 2C). Endosymbionts cannot freely produce biomass, but must convert other metabolites into biomass. This is expressed by the second equality, which states that the net production of metabolites other than biomass 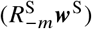 must be balanced by the net uptake of metabolites from the host (***f***^S^), which is bounded by the uptake limits set by the host (***c***^H^) (Fig. 2C). To express the irreversibility of the reactions, we constrained the weights of the regulatory modes to be nonnegative (***w***^**S**^ ≥ **0**). Formulating endosymbiont metabolism as a maximization problem (Eq. 1) expresses the autonomy of endosymbiont growth. The maximization can be interpreted as combining physiological adaptation of the endosymbiont to the host body with microevolution within it, in which the fastest-growing mutants outcompete the others. The metabolic exchange at optimal endosymbiont growth is written as ***f*** ^S∗^(***c***^H^; *R*^S^), where the asterisk indicates optimality. At this optimum, some metabolites are consumed up to their host-provided limit and others are not. We refer to the former as bounded and the latter as unbounded (Fig. 2C). This distinction determines how endosymbionts respond to their hosts: changing the uptake limit of a bounded metabolite alters endosymbiont metabolism, whereas changing that of an unbounded metabolite does not, because it is already in excess.
2. **Host adaptation:** Next, the host adapts to its endosymbiont by maximizing its growth while taking up metabolites from the environment at rates up to ***c***^E^, given the metabolites that its endosymbiont consumes or excretes (***f*** ^S∗^).

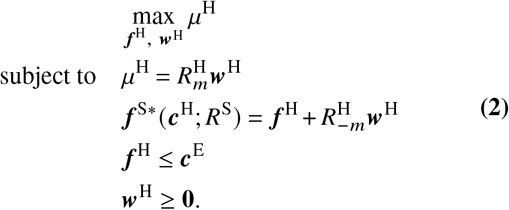

To model host dependence on endosymbionts, we selected, among hosts with randomly generated regulatory matrices *R*^H^, those that grow poorly without their endosymbionts in an environment where only the first metabolite can be taken up (***c***^E^ = (1, 0, …, 0)^⊤^): their endosymbiont-free growth was low 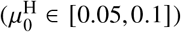 relative to the maximum possible growth (µ^H^ = 1) of a host able to use all available metabolites.

### The number of regulatory modes affects host– endosymbiont competition

Before explaining the third level of optimization (host evolution), we first examined how the number of endosymbiont regulatory modes (*r*) affects the metabolic competition between hosts and endosymbionts. We generated 300 endosymbionts with different random *R*^S^ for each *r* ranging from 1 to 100 (*N* = 30,000 total endosymbionts) and assigned them to hosts with the same *R*^H^ but varying internal environments (***c***^H^). We assigned random uptake limits ***c***^H^ to the non-biomass metabolites (total uptake limit ∥***c***^H^ ∥_1_ = 0.01≪ 1, with 100 random compositions per endosymbiont). Performing the first two levels of optimization (endosymbiont and host adaptation without host evolution) revealed clear effects of *r* (Fig. 3).

**Figure 3.**
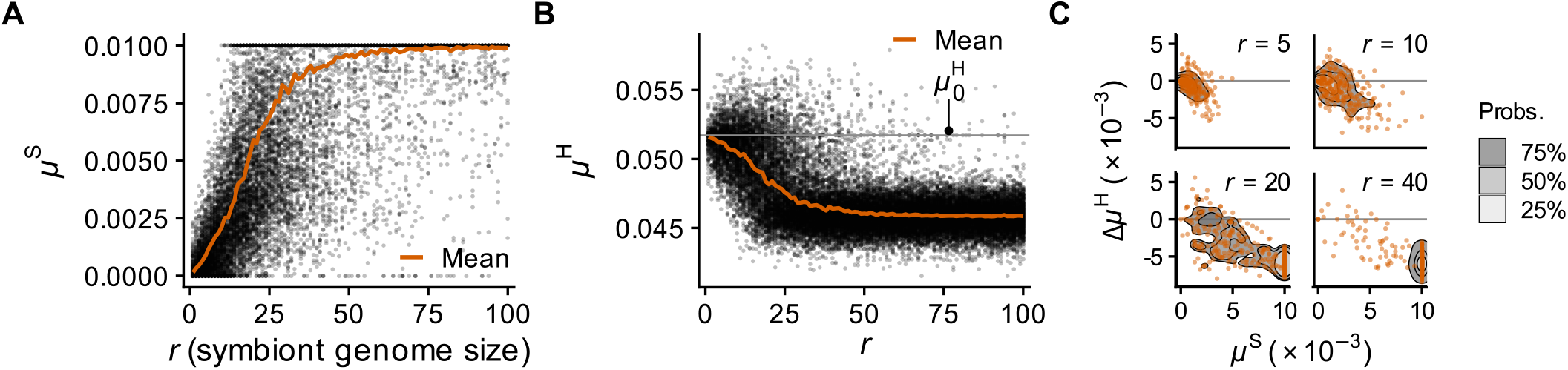
Metabolic competition between hosts and endosymbionts before host-level evolution. **(A, B)** Random endosymbionts with different numbers of regulatory modes (*r*) were assigned to hosts with varying internal environments (***c***^H^) but fixed *R*^H^. Each point corresponds to the growth (µ) of a host–endosymbiont pair with random ***c***^H^ and *R*^S^. Among the 100 random ***c***^H^ and 300 random *R*^S^ per *r*, only 300/30,000 random points per *r* are displayed for clarity. The thick orange lines show the means for each *r*, and the gray horizontal line shows the endosymbiont-free growth of the host 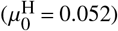. **(C)** The growth of host–endosymbiont pairs (points randomly subsampled to *n* = 300 per *r*). The contours show the highest-density regions, determined via kernel density estimation with a bandwidth of 2 × 10^−3^.

As expected, endosymbiont growth *µ*^S^ generally decreased as *r* decreased (Fig. 3A), reflecting the reduced metabolic capacity of endosymbionts with fewer regulatory modes to adapt to the host’s internal environment. Conversely, host growth µ^H^ increased on average as *r* decreased (Fig. 3B).

This inverse relationship reflects competition for shared resources, resulting in a negative correlation between host and endosymbiont growth (Spearman’s *ρ* = −0.58).

The competitive effects are clearly visible across the range of *r* (Fig. 3C). When *r* was large (*r* = 40), endosymbionts grew well at the expense of host resources, leading to lower host growth. As *r* decreased, endosymbionts became less capable of utilizing host-provided metabolites, allowing hosts to retain more resources and achieve higher growth. The parasitic burden imposed by endosymbionts was therefore smaller for endosymbionts with fewer regulatory modes, indicating that the loss of regulatory capacity reduces the immediate cost of endosymbiont acquisition.

These results demonstrate the baseline competitive relationships that hosts must overcome through evolution. The variation in responses across different host internal environments (***c***^H^) also provides the basis for calculating the sensitivity of endosymbiont metabolism, which we later use as an indicator of mutualistic potential.

### Mutualism evolved in hosts harboring endosymbionts with moderately reduced numbers of regulatory modes

**(3) Host evolution:** On top of levels (1) and (2), we incorporated the third level of optimization. Here, to model how hosts evolve to exploit their endosymbionts, we assume that hosts evolve to maximize their growth µ^H^ by tuning the uptake limits they impose on their endosymbionts (***c***^H^) inside their bodies (Fig. 2D). We assume that a host with a fixed *R*^H^ harboring an endosymbiont with a fixed *R*^S^ evolves to maximize its fitness µ^H^ by tuning ***c***^H^, which was held constant in the previous two levels of optimization:

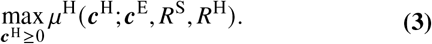

When host-level evolution was incorporated, hosts that acquired endosymbionts with moderately small numbers of regulatory modes *r* tended to achieve high fitness µ^H^, with a peak at *r*^∗^ = 13 (95% bootstrap CI 10–15; Fig. 4A). Hosts that acquired endosymbionts with large *r* (*r* > 25) evolved to suppress endosymbiont growth (*µ*^S^), preventing excessive consumption of metabolites that the host could otherwise utilize (Fig. 4B). In contrast, hosts with endosymbionts having smaller *r* (*r* < 25) allowed their endosymbionts to maintain higher growth rates. For instance, while at *r* = 40 both *µ*^S^ and µ^H^ were low, at *r* = 10, many host–endosymbiont pairs achieved high growth for both partners (Fig. 4C). We regarded a pair as mutualistic when, after host evolution, the host grew above its endosymbiont-free growth 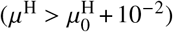 and the endosymbiont also maintained growth (*µ*^S^ >10^−2^). The same peak appeared when the analysis was restricted to the 24,787 of 30,000 pairs whose endosymbionts were parasitic before host evolution, that is, for which none of the 100 random ***c***^H^ raised host growth above 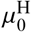 (*SI Appendix*, Fig. S1).

**Figure 4.**
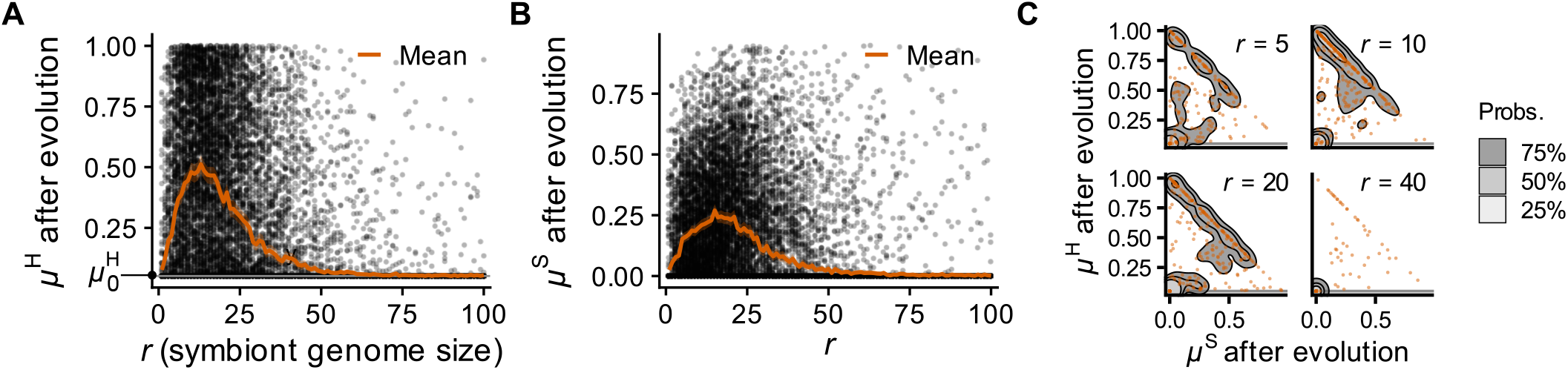
Evolution of mutualism after host-level evolution. **(A, B)** Random endosymbionts with varying genome sizes (*r*) were each assigned to a host with a fixed *R*^H^, and each host evolved to maximize its growth (µ^H^) by tuning its internal environment (***c***^H^) to exploit its endosymbiont. Each point corresponds to the growth (µ) of a random host–endosymbiont pair with fixed *R*^H^ and random *R*^S^ (*N* = 300 per *r*). **(C)** The growth of host–endosymbiont pairs (*N* = 300 per *r*). The contours show the highest-density regions, determined via kernel density estimation with a bandwidth of 0.2. The gray horizontal lines in A and C indicate endosymbiont-free growth 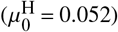.

This peak was not specific to the particular host regulatory matrix *R*^H^ used in the main analysis. It was consistent across different host regulatory matrices *R*^H^ (*SI Appendix*, Figs. S2A and S3). It also did not depend on the coupling between *r* and metabolic capacity: because each regulatory mode was assigned *k* = 4 reactions, the total number of reactions available to endosymbionts increases with *r*, particularly at small *r* (*SI*

*Appendix*, Fig. S2B). Varying *k* (*SI Appendix*, Fig. S2C), or fixing the total set of reactions while varying only its partition into regulatory modes (*SI Appendix*, Fig. S2D), changed the overall level of host growth but left the peak at intermediate *r* (*r*^∗^ = 8–16 across these scenarios). While evolution in theory can reach optima, in practice it may be constrained by the host’s ability to explore the space of possible ***c***^H^. Nevertheless, a genetic algorithm, which does not rely on the mixed-integer linear programming (MILP) solver that directly computes the optimal ***c***^H^, reached a host growth rate no more than 10^−2^ below that of the MILP solver for 98% of host–endosymbiont pairs, and yielded the same dependence of µ^H^ on *r*, with the maximum at the same *r* (*SI Appendix*, Fig. S4).

### The “adaptability” and “sensitivity” of endosymbionts

Host fitness (µ^H^) after evolution was strongly correlated with the maximum µ^H^ before evolution (*ρ* = 0.76 for 5≤ *r* ≤25; Fig. 5A) and less strongly with the median (*ρ* = 0.53). The correlation with the maximum was not explained by their common dependence on *r* (partial *ρ* = 0.78 controlling for *r*). This suggests that understanding the source of variation in pre-evolution host growth could reveal the mechanism underlying mutualism evolution at intermediate genome sizes.

**Figure 5.**
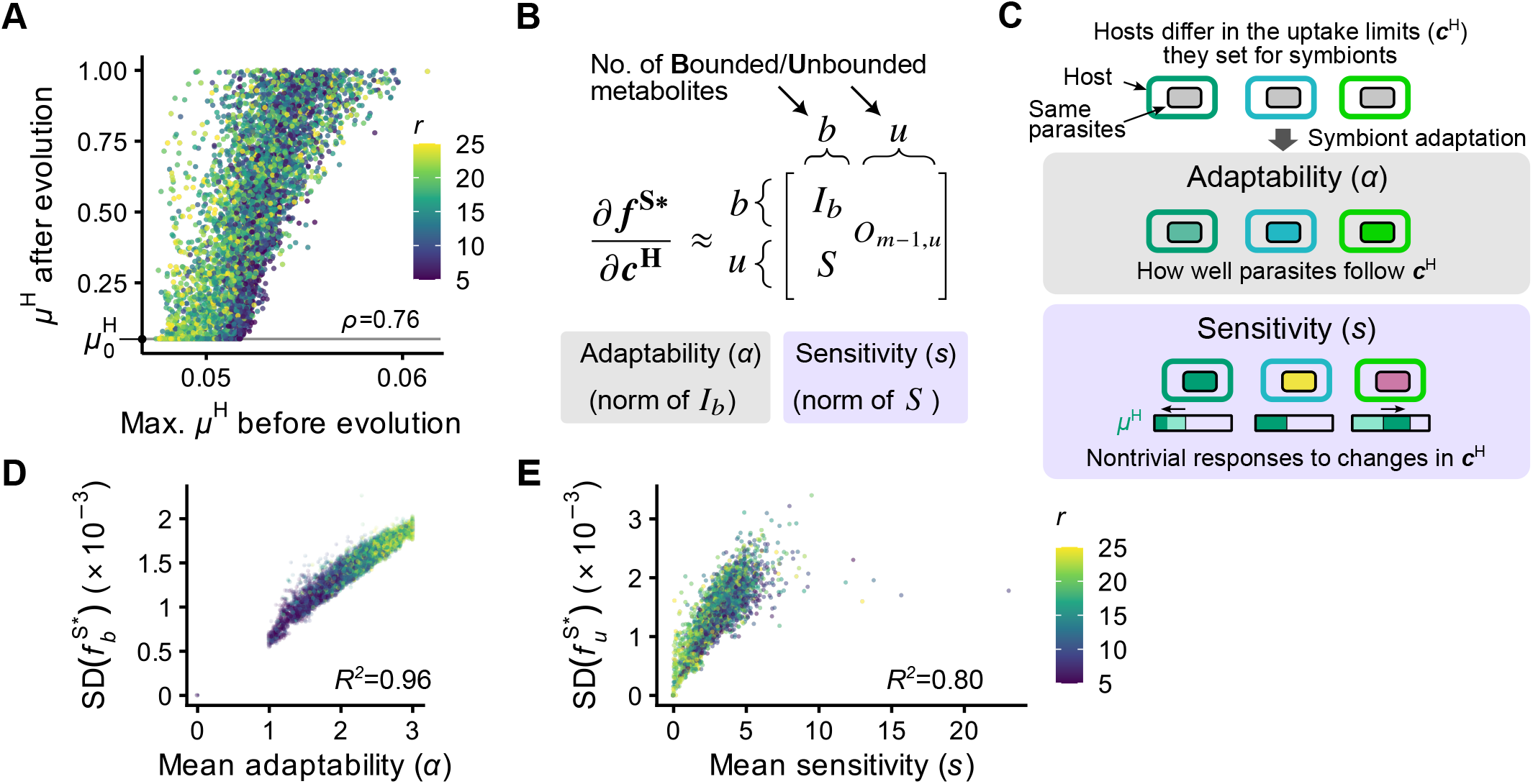
The decomposition of the response of endosymbiont metabolism to variation in host-provided metabolites. **(A)** Host fitness (µ^H^) after evolution (Fig. 4A) was strongly correlated with the maximum µ^H^ before evolution (Fig. 3B). **(B)** The response of metabolic exchange between host and endosymbiont (***f*** ^S∗^) to variation in host-provided metabolites (***c***^H^) can be decomposed into two components, the *adaptability* (α) and *sensitivity* (*s*) of endosymbiont metabolism, corresponding to the variation in the bounded (used up by the endosymbiont) and unbounded non-biomass metabolites, respectively. The rows and columns of the matrix are reordered such that the first *b* elements correspond to bounded metabolites that are completely utilized by the endosymbiont, and the remaining *u* elements correspond to unbounded metabolites. *I*_*b*_ denotes a *b* ×*b* identity matrix, and *O*_*m*−1,*u*_ denotes an (*m*_−_1) ×*u* zero matrix. **(C)** Schematic representation of the adaptability and sensitivity of endosymbiont metabolism. The two metrics capture different aspects of the responses of endosymbiont metabolism (fill colors of endosymbionts) to differences in host-provided metabolites (***c***^H^) (border colors of hosts). **(D, E)** The adaptability (α) and sensitivity (*s*) of endosymbiont metabolism scale approximately linearly with the variations in bounded 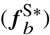 nand unbounded 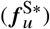 components of metabolic exchanges, respectively, across different host–endosymbiont pairs with endosymbiont genome sizes between 5 and 25 (*N* = 300 per *r*) as in Fig. 3. Each point shows the mean values of an endosymbiont with a specific regulatory matrix *R*^S^, experiencing different host-provided metabolites ***c***^H^.

To identify this mechanism, we analyzed how the metabolism of endosymbionts responds to variation in host-provided metabolites (***c***^H^). Since the environment (***c***^E^) and host regulatory matrix (*R*^H^) were held constant, changes in host growth (µ^H^) following changes in host-provided metabolites (***c***^H^) must result from altered metabolic exchanges with endosymbionts (***f*** ^S∗^). Therefore, we sought to quantify how changes in host-provided metabolites (***c***^H^) affect metabolic exchanges (***f*** ^S∗^) between the host and its endosymbionts, which can be expressed as the Jacobian matrix ∂ ***f*** ^S∗^/∂***c***^H^.

However, the nested optimization structure prevents direct computation of the Jacobian matrix. To overcome this, we made a local linear approximation that allows the decomposition of the Jacobian matrix into components related to the bounded and unbounded metabolites, or those that are consumed by the endosymbiont up to the maximum available in the host and those that are not, respectively (Fig. 2C) (see *SI Appendix*, section S7 for details). This approximation allows endosymbiont responses to be decomposed into two components (Fig. 5BC): **adaptability (**α): the capacity to directly respond to changes in host-provided metabolites, and **sensitivity (***s*): the propensity for indirect metabolic responses with the potential to benefit hosts.

Under this approximation, the Jacobian takes the block form (Fig. 5B):

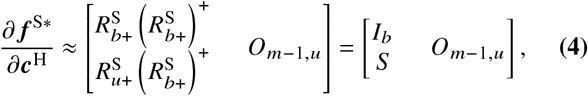

where the rows and columns are ordered so that the first *b* correspond to the bounded metabolites and the remaining *u* to the unbounded ones. 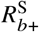 and 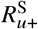 are submatrices of *R*^S^ : the subscripts *b* and *u* select the rows of bounded and unbounded metabolites, respectively, and the subscript selects the columns of the regulatory modes with nonzero weights in the optimal endosymbiont metabolism. 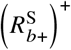 is the Moore– Penrose pseudoinverse of 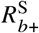. The upper block equals the *b* × *b* identity matrix *I*_*b*_ because 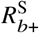 has full row rank at almost every optimum (*SI Appendix*, section S7), and *O*_*m*−1,*u*_ is a zero matrix because the endosymbiont does not respond to metabolites already in excess.

We define the adaptability and the sensitivity of the endosymbiont as the norms of the upper and lower blocks, respectively:

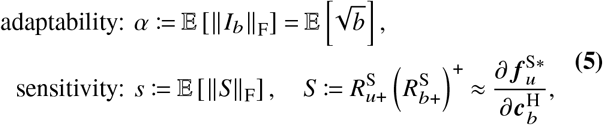

where 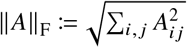 is the Frobenius norm of matrix *A*, and E[·] denotes the average over different compositions of metabolites supplied by the host, ***c***^H^. The sensitivity matrix *S* captures the indirect metabolic responses of the endosymbiont to host-provided metabolites, in contrast to the identity matrix *I*_*b*_, which captures the direct responses.

Each component captures a different aspect of the endosymbiont’s metabolic response to host-provided metabolites (Fig. 5BC). Adaptability (α) measures how changes in the bounded host-provided metabolites 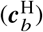 affect the bounded metabolic exchanges 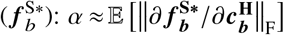. In contrast, sensitivity (*s*) measures how changes in 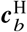 affect the unbounded metabolic exchanges 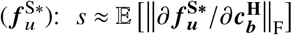. In other words, adaptability (α) captures how well endosymbionts can follow changes in the available host-provided metabolites 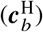, whereas sensitivity (*s*) captures indirect responses that do not simply reflect consumption of all available host-provided resources. These indirect responses are nontrivial, as captured by the structure of the sensitivity matrix *S*, and can produce metabolic outputs that benefit the host.

This interpretation is consistent with the observed correlations between these components and the variation in metabolic exchanges. Intuitively, under a local linear-response approximation, variation in metabolic exchanges satisfies 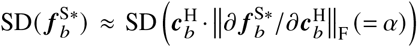. Thus, when α is approximately constant over the range of host-provided metabolite profiles considered, 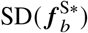 is expected to scale with adaptability (α). Indeed, adaptability (α) scaled approximately linearly with 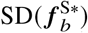 (ordinary least-squares fit: *R*^2^ = 0.96, intercept: 8.8 × 10^−^; Fig. 5D) for the small ∥***c***^H^ ∥_1_ used in Fig. 3B (see *SI Appendix*, section S9.1 for the definition of SD). Similarly, sensitivity scaled approximately linearly with 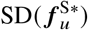 (*R*^2^ = 0.80, intercept: 2.7 × 10^−4^; Fig. 5E), consistent with the local linear approximation.

### Sensitivity requires metabolites the host can control and metabolites it can use

We next asked why hosts exploited endosymbionts most readily at intermediate *r* (Fig. 4A). The decomposition above suggests that this reflects two things hosts need from their endosymbionts. First, hosts must be able to alter endosymbiont metabolism, and second, those changes must be able to benefit the host. Bounded metabolites (*b*) enable host control: because the endosymbiont takes them up to their limits, the host can manipulate their uptake by changing these limits. Unbounded metabolites (*u*) provide the benefit: their fluxes are not pinned at the uptake limits, so they can be excreted in forms the host can use. Mutualism thus requires both *b* > 0 and *u* > 0, which is exactly when the sensitivity matrix *S* is nonempty.

However, *b* and *u* cannot both be large. Because *b* +*u* = *n* is fixed as the total number of non-biomass metabolites (*n* = *m* −1) while *b* increases with *r* (Fig. 6A), endosymbionts with few regulatory modes give the host little to control, whereas those with many exhaust nearly every metabolite and leave little to excrete. The product *ub*, which is the number of elements of *S* (Eq. 5), is largest at *b* = *n* /2, which the mean *b* reaches at *r* ≈12 (Fig. 6A). We therefore expect sensitivity, and with it the potential for mutualism, to peak at intermediate *r*, and the position of this peak to depend on the number of metabolites.

**Figure 6.**
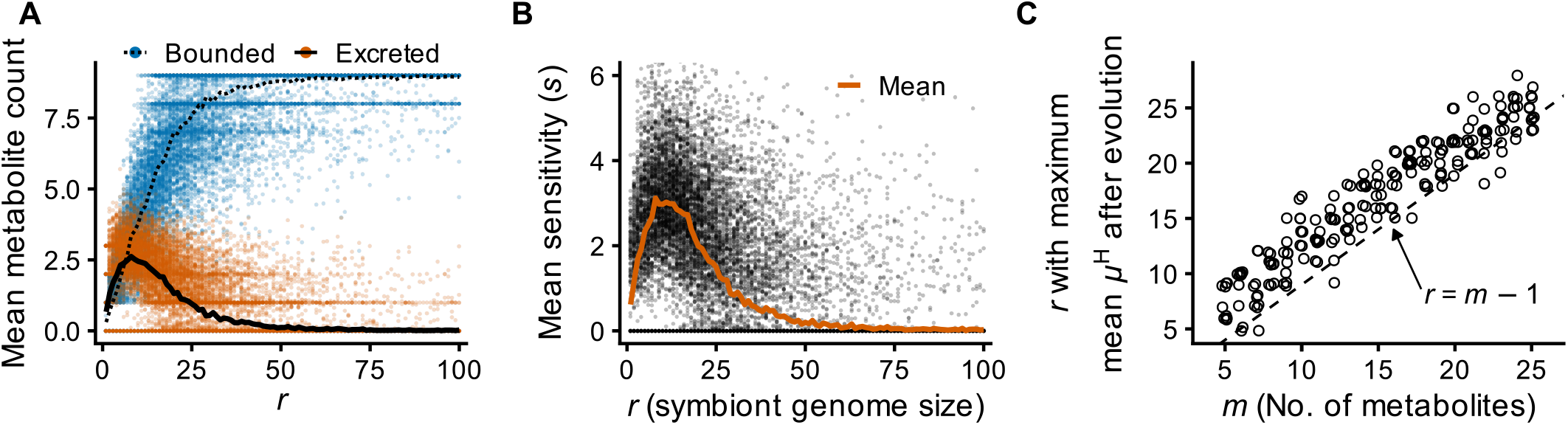
Endosymbiont sensitivity and the *r* that maximized host growth after evolution. **(A)** The mean numbers of bounded metabolites (blue) and of excreted metabolites (orange) per endosymbiont before host evolution. **(B)** The sensitivity of endosymbionts (*s*) peaked at intermediate genome sizes (*N* = 300 per *r*). **(C)** The *r* that achieved the maximum mean host growth rate µ^H^ after evolution increased linearly with *m*. Each point indicates the result from the evolution of a host with a randomly generated *R*^H^ (*r*^H^ = 24; *N* = 10 hosts per *m* from 5 to 25). For *m* = 10, each host was evolved with 300 random endosymbionts per genome size (*r* = 1, 2, …, 100) as in Fig. 4A; for other *m*, with 100 random endosymbionts at each of 18–22 genome sizes between *r* = 1 and 100, sampled more densely around *r* = *m* −1. The mean µ^H^ after evolution for each endosymbiont genome size (*r*) was calculated, and the *r* that achieved maximum mean µ^H^ after evolution is shown. The dashed line shows *r* = *m* − 1. The points are jittered by at most 0.2 on both axes to improve visibility.

Sensitivity indeed peaked at intermediate *r* (Fig. 6B), similar to the peak in host growth µ^H^ after evolution (Fig. 4A). Nonzero sensitivity was nearly a prerequisite for mutualism: only 0.3% of pairs with *s* = 0 became mutualistic, whereas 99% of mutualistic pairs had *s* > 0 (*SI Appendix*, Fig. S5A), and the fraction of endosymbionts with *s* > 0 peaked at the same *r* as µ^H^ after evolution (*SI Appendix*, Fig. S5C). At a given *r*, higher sensitivity was associated with both a larger fraction of pairs becoming mutualistic and higher host growth in those pairs (*SI Appendix*, Fig. S5AB), although only modestly, likely because *s* measures the magnitude of these responses but not whether they yield metabolites the host can use, which depends on the host’s own regulatory network and environment.

The peak position also changed with the number of metabolites as expected: the *r* that maximizes µ^H^ after evolution lies slightly above *n* for *m* from 5 to 25 (Fig. 6C, *SI Appendix*, Fig. S6), which is consistent with an argument based on the number of tight constraints in the endosymbiont optimization problem (*SI Appendix*, section S8).

### Endosymbiont genome size evolved toward the sensitivity peak

While the above results show that hosts can exploit endosymbionts with intermediate genome sizes, they do not show that evolution leads to this outcome. We therefore added a fourth level on top of the three: the evolution of endosymbiont regulatory matrices, in which endosymbionts altered their regulatory matrices *R*^S^ through random gains and losses of regulatory modes and faster-growing hosts were preferentially selected (*SI Appendix*, section S5.3).

Endosymbionts starting from *r*_0_ = 5 gained regulatory modes and those starting from *r*_0_ = 30 and above lost them, so that all populations moved toward the region in which µ^H^ after host evolution is highest (Fig. 7). They did not, however, converge on a single genome size: individual lineages wandered widely, reaching *r* > 50 even when they had started at *r*_0_ = 5 (*SI Appendix*, Fig. S7). This variation is consistent with selection on *r* being weak, as *r* affects host fitness only indirectly, through how far the host can evolve to exploit its endosymbiont. Therefore, while selection favors intermediate genome sizes, it is likely not strong enough to prevent endosymbionts from wandering to larger or smaller genome sizes.

**Figure 7.**
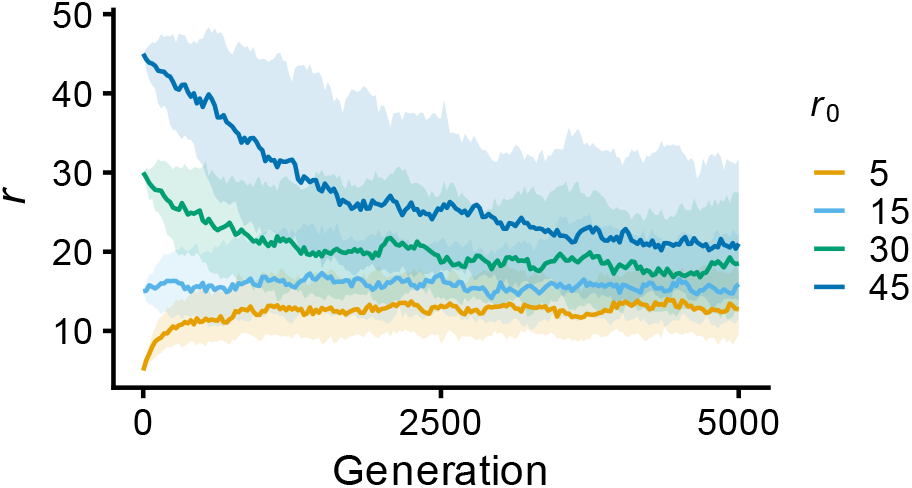
Evolution of the number of regul0atory modes (*r*). Endosymbionts altered their regulatory matrices *R*^S^ through random gains and losses of regulatory modes, starting from *r*_0_ = 5, 15, 30 and 45 (160 replicate populations per *r*_0_). Lines show the median *r* across replicates and shaded regions the interquartile range.

## Discussion

Endosymbionts typically undergo extreme genome reduction (McCutcheon and Moran, 2012), but how this affects the ability of hosts to exploit them has remained unclear. Using a model of metabolic competition between hosts and endosymbionts (Fig. 2), we found that hosts most readily evolve to exploit endosymbionts of intermediate genome size (Fig. 4). Decomposing the endosymbiont’s response to host-provided metabolites into adaptability and sensitivity (Fig. 5) showed that sensitivity peaks at the same intermediate sizes and is nearly a prerequisite for mutualism (Fig. 6B and *SI Appendix*, Fig. S5). Endosymbionts must therefore be neither too autonomous nor too degraded, retaining enough regulatory capacity to respond to the host while producing metabolites useful to it.

Genome reduction is commonly viewed as passive decay under relaxed selection (O’Fallon, 2008; McCutcheon and Moran, 2012). Genome-reduced endosymbionts have elevated *dN*/ *dS* and accumulate mobile genetic elements such as insertion sequences (McCutcheon and Moran, 2012), both of which are taken as evidence of neutral evolution. Our results suggest that these patterns may also reflect host-level selection: disruption of regulatory genes by nonsynonymous mutations and insertions, and the formation of new operons by insertion sequences (Kanai et al., 2022), remove or merge regulatory modes and thereby move an endosymbiont toward the sensitivity peak. Host control has already been implicated in the loss of symbiont traits, such as flagella in host-associated bacteria (Sharp and Foster, 2022).

In our model, what makes endosymbionts exploitable is the number of independently regulated units rather than their metabolic content, which has been the focus of much work on endosymbiont genome reduction (McCutcheon and Moran, 2012). Reducing *r* in our main analyses removes both, but the peak in host growth persisted when the metabolic repertoire was held fixed and only its partition into regulatory modes varied (*SI Appendix*, Fig. S2D). In real genomes, regulatory genes scale more steeply with genome size than metabolic genes (Fig. 1A; (Galán-Vásquez et al., 2016)), and the ratio of regulatory to metabolic genes declines with decreasing genome size even within lineages containing intermediate-stage endosymbionts (Fig. 1B). This is consistent with the view that mutualism can emerge through changes in gene regulation rather than through the acquisition of new genes (Shigenobu et al., 2000; Moran, 2003).

Our results suggest a trajectory from free-living autonomy to mutualism. Endosymbionts with large genomes retain autonomy and consume metabolites the host could otherwise use (Figs. 3C and 4A). As reduction proceeds, they pass through a sensitive phase (*r* ~*m*) in which hosts can select for beneficial metabolic exchanges, as may be the case for some *Arsenophonus, Sodalis* and *Serratia* endosymbionts (Boyd et al., 2024; Siozios et al., 2024; Manzano-Marín and Latorre, 2016). Extremely reduced endosymbionts become less responsive to host control (Fig. 6B, *r* < 5). Such endosymbionts, like *Buchnera*, have long coadapted to supply their hosts directly, and so no longer fit our assumption of autonomy. When the host faces a new environment, however, this loss of sensitivity may constrain host adaptation and help explain the frequent replacement of endosymbionts observed in nature (Henry et al., 2013; Husnik and McCutcheon, 2016; Sudakaran et al., 2017; Dial et al., 2022).

Theory on the evolution of mutualism has largely located its explanation in the structure of the interaction, such as transmission mode and the relative timescales of host and endosymbiont replication (Frank, 1996; Law and Dieckmann, 1998; van Vliet and Doebeli, 2019), or in the host’s capacity to choose, reward and punish partners (Bull and Rice, 1991; Kiers et al., 2003; Sachs et al., 2004). These frameworks explain how mutualism is reached and maintained but largely take variation in partner quality as given. Our mechanism adds a factor that lies in the endosymbiont’s genome. We assumed vertical transmission, and the host’s evolution consists only of adjusting the maximum rate at which the endosymbiont can take up each metabolite (Price et al., 2014; Wilde et al., 2024). What the endosymbiont returns remains a byproduct of its own growth maximization (Sachs et al., 2004), but one the host can steer by changing these rate limits (Fig. 5C). Our results suggest that genome reduction makes endosymbionts more susceptible to such steering. This mechanism does not exclude other routes, including loss of functions supplied by partners, whether through relaxed selection (Ellers et al., 2012) or adaptive gene loss (Morris et al., 2012), and selection to reduce the metabolic costs of symbiosis (Ankrah et al., 2018; Perreau and Moran, 2022). Genome reduction also occurs in pathogens such as *Mycoplasma* (Rocha and Blanchard, 2002), which likely have reduced genome sizes independently of host-level selection for mutualism.

Several aspects of the model could be improved in future studies. First, and most notably, we assumed that endosymbionts adapt much faster than hosts evolve, so that hosts always face endosymbionts at their optimum. While this assumption allowed us to derive the sensitivity of endosymbionts to host-provided metabolites, it is unlikely to hold in natural host–endosymbiont coevolution (Drew et al., 2021). When endosymbiont genotypes compete within hosts, selection favors faster-growing endosymbionts that retain regulatory capacity (Fig. 3A), opposing the host-level selection described here. Incorporating regulatory capacity into multilevel-selection models, in which the rate of horizontal transmission and the number of endosymbiont generations per host generation vary (Law and Dieckmann, 1998; van Vliet and Doebeli, 2019), would reveal how sensitivity-mediated host control interacts with transmission mode, with the evolution of host control over transmission itself (Frank, 1996), and with other forms of host control such as partner choice and sanctions (Kiers et al., 2003; Sachs et al., 2004). Such an analysis might also identify conditions under which endosymbionts reliably reach *r*^∗^, rather than wandering widely as in our model (Fig. 7). Second, our regulatory matrices are random, whereas real networks have structure that may affect sensitivity. Introducing realistic structures into the regulatory matrices (Klamt et al., 2017; Sastry et al., 2019), or using metabolic (Thomas et al., 2009; Ankrah et al., 2018) and gene regulatory networks (Yus et al., 2019) from real genome-reduced bacteria and their relatives, could identify genes or network motifs crucial for endosymbiont sensitivity. In particular, the loss of global regulators can shift bacteria from pathogenicity toward commensalism (Jansen et al., 2015) or even mutualism (Koga et al., 2022), possibly through abrupt changes in sensitivity. Whether global or local regulators are lost first during reduction could be examined in closely related lineages spanning different stages of reduction in nature (Hosokawa et al., 2016; Boscaro et al., 2017; Siozios et al., 2024; Boyd et al., 2024). Finally, sensitivity as defined here requires, in addition to *R*^S^, knowing which metabolites are bounded and which regulatory modes are active, information that is difficult to obtain in endosymbionts that cannot be cultured. To make sensitivity measurable, future studies could develop measures based instead on the genotype *R*^S^ alone, or on the activity of regulatory modes ***w***^S^. Transcriptomic responses to different media in *E. coli* strains with varying genome sizes (Kurokawa et al., 2016; Kanai et al., 2025) or in experimentally established endosymbioses (Mehta et al., 2018; Koga et al., 2022), together with the variation of endosymbiont gene expression with host genotype (Chong and Moran, 2016; Smith and Moran, 2020), would then make it possible to test whether sensitivity is indeed maximized at intermediate genome sizes.

Overall, our model offers a conceptual framework in which genome reduction is not mere degradation but an evolvability-enhancing process that facilitates the evolution of mutualism by increasing endosymbiont sensitivity to host-provided metabolites.

## Materials and Methods

At fixed ***c***^H^ (levels 1 and 2), the model reduces to two linear programs separated by a strictly convex quadratic program that selects the minimum-norm ***w***^S^ among the endosymbiont’s optima, which makes ***f*** ^S∗^ a single-valued function of (***c***^H^, *R*^S^). Host evolution (level 3) was solved by replacing the two lower levels with their Karush–Kuhn– Tucker conditions and imposing the complementarity constraints as type-1 special ordered sets, giving a mixed-integer linear program (Siddiqui and Gabriel, 2013). Problems that reached the solver time limit were represented by the best feasible solution found, which is a lower bound on the optimum; excluding them or bounding them from above did not change the position of the peak (*SI Appendix*, Fig. S8).

The Jacobian ∂ ***f*** ^S∗^/∂***c***^H^ on which adaptability and sensitivity are based (Eq. 5) cannot be computed directly through the nested optim izati on. We obtained it by approximating level 1 as 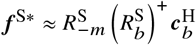, that is, as matching the endosymbiont’s turnover to the uptake limits of the metabolites it exhausts, and by restricting the derivative to the regulatory modes that are active at the optimum.

Reaction directions were drawn independently for every organism; sharing them between the host and its endosymbionts lowered the peak in µ^H^ but did not shift it (*SI Appendix*, Fig. S9). The generation of the regulatory matrices and of the internal environments, the numerical solution, the derivation of the sensitivity matrix, the quantities computed from the solutions, the genomic analysis and the statistical analysis are described in *SI Appendix*.

## Data, materials and software availability

Code used to generate and analyze the model will be made available on GitHub upon publication.

## Acknowledgments

The authors thank Dr. Saburo Tsuru for helpful comments on the manuscript, Dr. Nen Saito for helpful discussions on the model formulation, Dr. Shuji Shigenobu for helpful discussions on genomic analysis, and members of the ERATO Fukatsu Symbiotic Evolution Project, including Dr. Takema Fukatsu and Dr. Ryuichi Koga, for discussions on the biological implications of the results. Y.K. thanks Dr. Filip Husnik for hosting a research visit and members of his unit for discussions during the development of this project.

This work was supported by JSPS KAKENHI grant numbers 21J20693 and 25K18448 to Y.K., and 22K21344, 24H01798, 26H01831 and 26K02000 to C.F.; the Noda Institute for Scientific Research Young Investigator Research Grant to Y.K.; the Japan Science and Technology Agency PRESTO grant number JPMJPR2532 to Y.K.; and the Japan Science and Technology Agency ERATO grant number JPMJER1902 to C.F.

## Author contributions

Conceptualization, Y.K. and C.F.; Methodology, Y.K. and C.F.; Software, Y.K.; Formal analysis, Y.K.; Investigation, Y.K. and C.F.; Resources, C.F.; Data curation, Y.K.; Writing— original draft, Y.K.; Writing—review & editing, Y.K. and C.F.; Visualization, Y.K.; Supervision, C.F.; Funding acquisition, Y.K. and C.F.

## References

Ankrah, N. Y. D., Chouaia, B., and Douglas, A. E. The cost of metabolic interactions in symbioses between insects and bacteria with reduced genomes. mBio, 9(5):10.1128/mbio.01433–18, 2018. doi:10.1128/mbio.01433-18.

Boscaro, V., Kolisko, M., Felletti, M., Vannini, C., Lynn, D. H., and Keeling, P. J. Parallel genome reduction in symbionts descended from closely related free-living bacteria. Nat. Ecol. Evol., 1(8):1160–1167, 2017. doi:10.1038/s41559-017-0237-0.

Boyd, B. M., James, I., Johnson, K. P., Weiss, R. B., Bush, S. E., Clayton, D. H., and Dale, C. Stochasticity, determinism, and contingency shape genome evolution of endosymbiotic bacteria. Nat. Commun., 15:4571, 2024. doi:10.1038/s41467-024-48784-2.

Bull, J. J. and Rice, W. R. Distinguishing mechanisms for the evolution of co-operation. Journal of Theoretical Biology, 149(1):63–74, 1991. doi:10.1016/S0022-5193(05)80072-4.

Chong, R. A. and Moran, N. A. Intraspecific genetic variation in hosts affects regulation of obligate heritable symbionts. Proceedings of the National Academy of Sciences, 113(46):13114–13119, 2016. doi:10.1073/pnas.1610749113.

Cornwallis, C. K., van ‘t Padje, A., Ellers, J., Klein, M., Jackson, R., Kiers, E. T., West, S. A., and Henry, L. M. Symbioses shape feeding niches and diversification across insects. Nat. Ecol. Evol., 7(7):1022–1044, 2023. doi:10.1038/s41559-023-02058-0.

Cui, G., Liew, Y. J., Li, Y., Kharbatia, N., Zahran, N. I., Emwas, A.-H., Eguiluz, V. M., and Aranda, M. Host-dependent nitrogen recycling as a mechanism of symbiont control in Aiptasia. PLOS Genetics, 15(6):e1008189, 2019. doi:10.1371/journal.pgen.1008189.

Dial, D. T., Weglarz, K. M., Aremu, A. O., Havill, N. P., Pearson, T. A., Burke, G. R., and von Dohlen, C. D. Transitional genomes and nutritional role reversals identified for dual symbionts of adelgids (Aphidoidea: Adelgidae). ISME J., 16(3):642–654, 2022. doi:10.1038/s41396-021-01102-w.

Drew, G. C., Stevens, E. J., and King, K. C. Microbial evolution and transitions along the parasite–mutualist continuum. Nat. Rev. Microbiol., 19(10):623–638, 2021. doi:10.1038/s41579-021-00550-7.

Ellers, J., Toby Kiers, E., Currie, C. R., McDonald, B. R., and Visser, B. Ecological interactions drive evolutionary loss of traits. Ecology Letters, 15 (10):1071–1082, 2012. doi:10.1111/j.1461-0248.2012.01830.x.

Forni, G., Martelossi, J., Morel, B., Pistone, D., Bandi, C., and Montagna, M. Large-scale phylogenomics reveals convergent genome evolution across repeated transitions to endosymbiosis in Enterobacterales. Molecular Phylogenetics and Evolution, 217:108532, 2026. doi:10.1016/j.ympev.2026.108532.

Frank, S. A. Host control of symbiont transmission: The separation of symbionts into germ and soma. Am. Nat., 148(6):1113–1124, 1996. doi:10.1086/285974.

Gabr, A., Stephens, T. G., and Bhattacharya, D. Loss of key endosymbiont genes may facilitate early host control of the chromatophore in Paulinella. iScience, 25(9):104974, 2022. doi:10.1016/j.isci.2022.104974.

Galán-Vásquez, E., Sánchez-Osorio, I., and Martínez-Antonio, A. Transcription factors exhibit differential conservation in bacteria with reduced genomes. PLOS ONE, 11(1):e0146901, 2016. doi:10.1371/journal.pone.0146901.

Henry, L. M., Peccoud, J., Simon, J.-C., Hadfield, J. D., Maiden, M. J. C., Ferrari, J., and Godfray, H. C. J. Horizontally transmitted symbionts and host colonization of ecological niches. Curr. Biol., 23(17):1713–1717, 2013. doi:10.1016/j.cub.2013.07.029.

Hosokawa, T., Ishii, Y., Nikoh, N., Fujie, M., Satoh, N., and Fukatsu, T. Obligate bacterial mutualists evolving from environmental bacteria in natural insect populations. Nature Microbiology, 1:15011, 2016. doi:10.1038/nmicrobiol.2015.11.

Husnik, F. and McCutcheon, J. P. Repeated replacement of an intrabacterial symbiont in the tripartite nested mealybug symbiosis. Proceedings of the National Academy of Sciences, 113(37):E5416–E5424, 2016. doi:10.1073/pnas.1603910113.

Jansen, G., Crummenerl, L. L., Gilbert, F., Mohr, T., Pfefferkorn, R., Thänert, R., Rosenstiel, P., and Schulenburg, H. Evolutionary transition from pathogenicity to commensalism: Global regulator mutations mediate fitness gains through virulence attenuation. Mol. Biol. Evol., 32(11):2883–2896, 2015. doi:10.1093/molbev/msv160.

Kanai, Y., Tsuru, S., and Furusawa, C. Experimental demonstration of operon formation catalyzed by insertion sequence. Nucleic Acids Res., 50(3):1673–1686, 2022. doi:10.1093/nar/gkac004.

Kanai, Y., Shibai, A., Yokoi, N., Tsuru, S., and Furusawa, C. Laboratory evolution of the bacterial genome structure through insertion sequence activation. Nucleic Acids Res., 53(9):gkaf331, 2025. doi:10.1093/nar/gkaf331.

Keeling, P. J. and McCutcheon, J. P. Endosymbiosis: The feeling is not mutual. J. Theor. Biol., 434:75–79, 2017. doi:10.1016/j.jtbi.2017.06.008.

Kiers, E. T., Rousseau, R. A., West, S. A., and Denison, R. F. Host sanctions and the legume–rhizobium mutualism. Nature, 425(6953):78–81, 2003. doi:10.1038/nature01931.

Klamt, S., Regensburger, G., Gerstl, M. P., Jungreuthmayer, C., Schuster, S., Mahadevan, R., Zanghellini, J., and Müller, S. From elementary flux modes to elementary flux vectors: Metabolic pathway analysis with arbitrary linear flux constraints. PLOS Comput. Biol., 13(4):e1005409, 2017. doi:10.1371/journal.pcbi.1005409.

Koga, R., Moriyama, M., Onodera-Tanifuji, N., Ishii, Y., Takai, H., Mizutani, M., Oguchi, K., Okura, R., Suzuki, S., Gotoh, Y., Hayashi, T., Seki, M., Suzuki, Y., Nishide, Y., Hosokawa, T., Wakamoto, Y., Furusawa, C., and Fukatsu, T. Single mutation makes Escherichia coli an insect mutualist. Nat. Microbiol., 7(8):1141–1150, 2022. doi:10.1038/s41564-022-01179-9.

Kurokawa, M., Seno, S., Matsuda, H., and Ying, B.-W. Correlation between genome reduction and bacterial growth. DNA Research, 23(6):517–525, 2016. doi:10.1093/dnares/dsw035.

Larsson, J., Nylander, J. A., and Bergman, B. Genome fluctuations in cyanobacteria reflect evolutionary, developmental and adaptive traits. BMC Evolutionary Biology, 11(1):187, 2011. doi:10.1186/1471-2148-11-187.

Law, R. and Dieckmann, U. Symbiosis through exploitation and the merger of lineages in evolution. Proc. R. Soc. Lond. B Biol. Sci., 265(1402):1245–1253, 1998. doi:10.1098/rspb.1998.0426.

Lhee, D., Ha, J.-S., Kim, S., Park, M. G., Bhattacharya, D., and Yoon, H. S. Evolutionary dynamics of the chromatophore genome in three photosynthetic Paulinella species. Scientific Reports, 9(1):2560, 2019. doi:10.1038/s41598-019-38621-8.

Manzano-Marín, A. and Latorre, A. Snapshots of a shrinking partner: Genome reduction in Serratia symbiotica. Sci. Rep., 6(1):32590, 2016. doi:10.1038/srep32590.

McCutcheon, J. P. and Moran, N. A. Extreme genome reduction in symbiotic bacteria. Nat. Rev. Microbiol., 10:13–26, 2012. doi:10.1038/nrmicro2670.

Mehta, A. P., Supekova, L., Chen, J.-H., Pestonjamasp, K., Webster, P., Ko, Y., Henderson, S. C., McDermott, G., Supek, F., and Schultz, P. G. Engineering yeast endosymbionts as a step toward the evolution of mitochondria. Proc. Natl. Acad. Sci. U. S. A., 115(46):11796–11801, 2018. doi:10.1073/pnas.1813143115.

Mira, A., Ochman, H., and Moran, N. A. Deletional bias and the evolution of bacterial genomes. Trends Genet., 17(10):589–596, 2001. doi:10.1016/S0168-9525(01)02447-7.

Moran, N. A. Tracing the evolution of gene loss in obligate bacterial symbionts. Curr. Opin. Microbiol., 6(5):512–518, 2003. doi:10.1016/j.mib.2003.08.001.

Morris, J. J., Lenski, R. E., and Zinser, E. R. The black queen hypothesis: Evolution of dependencies through adaptive gene loss. mBio, 3(2): 10.1128/mbio.00036–12, 2012. doi:10.1128/mbio.00036-12.

Nechitaylo, T. Y., Sandoval-Calderón, M., Engl, T., Wielsch, N., Dunn, D. M., Goesmann, A., Strohm, E., Svatoš, A., Dale, C., Weiss, R. B., and Kaltenpoth, M. Incipient genome erosion and metabolic streamlining for antibiotic production in a defensive symbiont. Proc. Natl. Acad. Sci. U. S. A., 118(17):e2023047118, 2021. doi:10.1073/pnas.2023047118.

Nowack, E. C. M. and Melkonian, M. Endosymbiotic associations within protists. Philos. Trans. R. Soc. B Biol. Sci., 365(1541):699–712, 2010. doi:10.1098/rstb.2009.0188.

Nuñez, P. A., Romero, H., Farber, M. D., and Rocha, E. P. Natural selection for operons depends on genome size. Genome Biol. Evol., 5(11):2242–2254, 2013. doi:10.1093/gbe/evt174.

O’Fallon, B. Population structure, levels of selection, and the evolution of intracellular symbionts. Evolution, 62(2):361–373, 2008. doi:10.1111/j.1558-5646.2007.00289.x.

Pál, C., Papp, B., Lercher, M. J., Csermely, P., Oliver, S. G., and Hurst, L. D. Chance and necessity in the evolution of minimal metabolic networks. Nature, 440(7084):667–670, 2006. doi:10.1038/nature04568.

Perreau, J. and Moran, N. A. Genetic innovations in animal–microbe symbioses. Nat. Rev. Genet., 23(1):23–39, 2022. doi:10.1038/s41576-021-00395-z.

Prell, J., White, J. P., Bourdes, A., Bunnewell, S., Bongaerts, R. J., and Poole, P. S. Legumes regulate Rhizobium bacteroid development and persistence by the supply of branched-chain amino acids. Proceedings of the National Academy of Sciences, 106(30):12477–12482, 2009. doi:10.1073/pnas.0903653106.

Price, D. R. G., Duncan, R. P., Shigenobu, S., and Wilson, A. C. C. Genome Expansion and Differential Expression of Amino Acid Transporters at the Aphid/Buchnera Symbiotic Interface. Molecular Biology and Evolution, 28 (11):3113–3126, 2011. doi:10.1093/molbev/msr140.

Price, D. R. G., Feng, H., Baker, J. D., Bavan, S., Luetje, C. W., and Wilson, A. C. C. Aphid amino acid transporter regulates glutamine supply to intracellular bacterial symbionts. Proceedings of the National Academy of Sciences, 111(1):320–325, 2014. doi:10.1073/pnas.1306068111.

Rocha, E. P. C. and Blanchard, A. Genomic repeats, genome plasticity and the dynamics of Mycoplasma evolution. Nucleic Acids Res., 30(9):2031–2042, 2002. doi:10.1093/nar/30.9.2031.

Sachs, J. L., Mueller, U. G., Wilcox, T. P., and Bull, J. J. The evolution of cooperation. Q. Rev. Biol., 79(2):135–160, 2004. doi:10.1086/383541.

Sachs, J. L., Skophammer, R. G., Bansal, N., and Stajich, J. E. Evolutionary origins and diversification of proteobacterial mutualists. Proc. R. Soc. B Biol. Sci., 281(1775):20132146, 2014. doi:10.1098/rspb.2013.2146.

Sastry, A. V., Gao, Y., Szubin, R., Hefner, Y., Xu, S., Kim, D., Choudhary, K. S., Yang, L., King, Z. A., and Palsson, B. O. The Escherichia coli transcriptome mostly consists of independently regulated modules. Nat. Commun., 10(1):5536, 2019. doi:10.1038/s41467-019-13483-w.

Sela, I., Wolf, Y. I., and Koonin, E. V. Selection and genome plasticity as the key factors in the evolution of bacteria. Physical Review X, 9(3):031018, 2019. doi:10.1103/PhysRevX.9.031018.

Sharp, C. and Foster, K. R. Host control and the evolution of cooperation in host microbiomes. Nature Communications, 13(1):3567, 2022. doi:10.1038/s41467-022-30971-8.

Shigenobu, S., Watanabe, H., Hattori, M., Sakaki, Y., and Ishikawa, H. Genome sequence of the endocellular bacterial symbiont of aphids Buchnera sp. APS. Nature, 407(6800):81–86, 2000. doi:10.1038/35024074.

Siddiqui, S. and Gabriel, S. A. An SOS1-based approach for solving MPECs with a natural gas market application. Networks and Spatial Economics, 13 (2):205–227, 2013. doi:10.1007/s11067-012-9178-y.

Siozios, S., Nadal-Jimenez, P., Azagi, T., Sprong, H., Frost, C. L., Parratt, S. R., Taylor, G., Brettell, L., Liew, K. C., Croft, L., King, K. C., Brockhurst, aM. A., Hypša, V., Novakova, E., Darby, A. C., and Hurst, G. D. D. Genome dynamics across the evolutionary transition to endosymbiosis. Current Biology, 34(24):5659–5670.e7, 2024. doi:10.1016/j.cub.2024.10.044.

Smith, T. E. and Moran, N. A. Coordination of host and symbiont gene expression reveals a metabolic tug-of-war between aphids and Buchnera. Proc. Natl. Acad. Sci. U. S. A., 117(4):2113–2121, 2020. doi:10.1073/pnas.1916748117.

Sudakaran, S., Kost, C., and Kaltenpoth, M. Symbiont acquisition and replacement as a source of ecological innovation. Trends Microbiol., 25 (5):375–390, 2017. doi:10.1016/j.tim.2017.02.014.

Thomas, G. H., Zucker, J., Macdonald, S. J., Sorokin, A., Goryanin, I., and Douglas, A. E. A fragile metabolic network adapted for cooperation in the symbiotic bacterium Buchnera aphidicola. BMC Syst. Biol., 3:24, 2009. doi:10.1186/1752-0509-3-24.

van Nimwegen, E. Scaling laws in the functional content of genomes. Trends in Genetics, 19(9):479–484, 2003. doi:10.1016/S0168-9525(03)00203-8.

van Vliet, S. and Doebeli, M. The role of multilevel selection in host microbiome evolution. Proc. Natl. Acad. Sci. U. S. A., 116(41):20591–20597, 2019. doi:10.1073/pnas.1909790116.

Wilde, J., Slack, E., and Foster, K. R. Host control of the microbiome: Mechanisms, evolution, and disease. Science, 385(6706):eadi3338, 2024. doi:10.1126/science.adi3338.

Yus, E., Lloréns-Rico, V., Martínez, S., Gallo, C., Eilers, H., Blötz, C., Stülke, J., Lluch-Senar, M., and Serrano, L. Determination of the gene regulatory network of a genome-reduced bacterium highlights alternative regulation independent of transcription factors. Cell Syst., 9(2):143–158.e13, 2019. doi:10.1016/j.cels.2019.07.001.

