## Supplementary material for "Genome reduction increases endosymbiont sensitivity and promotes mutualism evolution": SI Appendix

### SI Appendix for

**This PDF file includes:**

- Supplementary Text
- Figs. S1 to S9
- Table S1
- SI References

### Supplementary Text

#### S1 Analyzing regulatory gene counts in genomes

Genomes from Enterobacterales (Forni et al., 2026), supplemented with additional *Arsenophonus* (Siozios et al., 2024) and *Serratia* genomes (Manzano-Marín and Latorre, 2016), were used to cover a broad range of genome sizes in clades that include major insect endosymbionts. These genomes comprised representatives of major insect-associated enterobacterial endosymbiont lineages, including *Arsenophonus*, *Baumannia*, *Blochmannia*, *Buchnera*, *Erwinia*, *Fukatsuia*, *Hamiltonella*, *Ishikawaella*, *Moranella*, *Mikella*, *Pantoea*, *Regiella*, *Riesia*, *Serratia*, *Sodalis*, *Symbiopectobacterium*, *Tachikawaea*, *Wigglesworthia* and *Westeberhardia*. RefSeq annotations generated by PGAP were used when available, and other RefSeq or GenBank annotations were used when PGAP-based annotations were unavailable. Genes were assigned to functional categories based on keywords in their product names. Genes were categorized as regulatory when their product names contained one of the following terms: "transcriptional regulator", "transcription factor", "transcriptional antiterminator", "DNA-binding", "sigma factor", "factor sigma", "response regulator", "transcriptional repressor", "transcriptional activator", "regulatory protein", "LysR", "TetR", "LuxR", "AraC", "GntR", or "MarR". Genes were categorized as metabolic when their product names contained one of the following terms: "synthase", "kinase", "dehydrogenase", "reductase", "transferase", "hydrolase", "oxidase", "lyase", "isomerase", "mutase", "glucosidase", or "deaminase".

Genomes of cyanobacteria (Larsson et al., 2011) and *Paulinella* chromatophores (Lhee et al., 2019) were included to cover a broad range of genome sizes among photosynthetic endosymbionts and their relatives. For cyanobacteria, genes from RefSeq annotations were used. For chromatophores, genes were predicted using GeneMarkS-2 (Lomsadze et al., 2018), which is also used in PGAP, to maintain consistency with RefSeq-based annotations. Genes from genomes of both origins were annotated using eggNOG-mapper v2 with the eggNOG 5.0 database (Cantalapiedra et al., 2021), in fast mapping mode, with the database limited to taxid 1117 (Cyanobacteriota). Genes assigned to COG category K were categorized as regulatory, whereas genes assigned to COG categories C, E, F, G, H, I, P, or Q were categorized as metabolic.

For both datasets, the total ungapped genome size was used as the genome size. The scaling relationship between the number of genes with specific functions (regulatory or metabolic) and genome size was analyzed by fitting the logarithm of the number of genes to the logarithm of the genome size using robust linear regression (`rlm` function, MASS package) in R.

#### S2 Generating random hosts, endosymbionts and internal environments

Hosts and endosymbionts were each assigned a regulatory matrix, $R^{\mathrm{H}}$ and $R^{\mathrm{S}}$, with $m$ rows corresponding to the metabolites and $r^{\mathrm{H}}$ and $r$ columns, respectively, corresponding to the regulatory modes. The matrices were generated in two steps. First, a reaction pool was drawn by assigning a direction uniformly at random to each of the ${}_m\mathrm{C}_2$ possible one-to-one conversions between the $m$ metabolites. This draw was made independently for every generated organism, so that hosts and endosymbionts share the set of metabolites but not the directions of their reactions.

Sharing reaction directions lowered the peak in $\mu^{\mathrm{H}}$ and made the outcome depend more on the reactions drawn, likely because the host and endosymbiont become more similar and thus more likely to compete for the same metabolites (Fig. S9A–E). Nevertheless, the peak in $\mu^{\mathrm{H}}$ at intermediate $r$, the $r$ that maximizes $\mu^{\mathrm{H}}$, and the peak in sensitivity were robust to sharing the reaction directions between hosts and endosymbionts (Fig. S9C–F).

Second, each regulatory mode was assembled by drawing $k$ reactions without replacement from that pool and assigning them weights drawn independently from $U(0,1)$; modes were drawn independently of one another, so the same reaction may appear in several modes.

We used $k = 4$ unless stated otherwise. This value was chosen so that many reactions are attached to each mode while two or more metabolites remain unconnected within a single mode ($4 \times 2 < m = 10$). Denser modes act on more metabolites at once, which restricts rather than enlarges the net uptakes the endosymbiont can realize and so leaves more metabolites unbounded. Larger $k$ therefore raises host growth at large $r$, but has little effect on the position of the peak (Fig. S2C).

Hosts were given $r^{\mathrm{H}} = 24$ regulatory modes unless stated otherwise, and endosymbionts $r$ from 1 to 100. To express the dependence of hosts on endosymbionts, randomly generated $R^{\mathrm{H}}$ were accepted only when the endosymbiont-free host growth in the environment $\boldsymbol{c}^{\mathrm{E}} = (1, 0, \ldots, 0)^\top$ satisfied $\mu_0^{\mathrm{H}} \in [0.05, 0.1]$, and rejected otherwise. The same $k$, the same $r^{\mathrm{H}}$ and the same acceptance window were used for the simulations with $m$ from 5 to 25. We also tested $r^{\mathrm{H}} = 2.4\,m$, which scales the host with the metabolic space it inhabits and coincides with $r^{\mathrm{H}} = 24$ at $m = 10$, with results qualitatively similar to Fig. 6C (Fig. S6).

The internal environments used before host evolution were generated by drawing $x_i \sim U(0,1)$ independently for the $m-1$ non-biomass metabolites and setting $\boldsymbol{c}^{\mathrm{H}} = 0.01\,\boldsymbol{x}/\sum_i x_i$, so that $\|\boldsymbol{c}^{\mathrm{H}}\|_1 = 0.01$. One hundred such environments were drawn per host–endosymbiont pair.

#### S3 The multilevel optimization of host–endosymbiont interaction

The phenotype of the host–endosymbiont pair was obtained through a set of maximization problems (Eqs. 1–3). Given $\boldsymbol{c}^{\mathrm{H}}$, $\boldsymbol{f}^{\mathrm{S}}$ is determined, which determines $\boldsymbol{\mu}^{\mathrm{H}}$, and the goal is to obtain $\boldsymbol{c}^{\mathrm{H}}$ that maximizes $\mu^{\mathrm{H}}$. Without additional constraints, there can be multiple solutions to the endosymbiont growth maximization problem (Eq. 1). When obtaining $\boldsymbol{c}^{\mathrm{H}}$, we added the following constraint to get a unique solution:

$$\min_{\boldsymbol{w}^{\mathrm{S}}} \|\boldsymbol{w}^{\mathrm{S}}\|_2^2 := \sum_{j=1}^{r} \left(w_j^{\mathrm{S}}\right)^2, \quad \text{s.t. } \mu^{\mathrm{S}} = \mu^{\mathrm{S}*}(c^{\mathrm{H}}; R^{\mathrm{S}}), \quad \text{(and constraints of Eq. 1)}, \tag{S1}$$

where $\mu^{\mathrm{S}*}$ is the optimal endosymbiont growth given $\boldsymbol{c}^{\mathrm{H}}$. This may be interpreted as endosymbionts minimizing the cost of gene expression.

To explicitly express the combined optimization problem, let us denote the set of combinations of $\boldsymbol{f}^{\mathrm{S}}$ and $\boldsymbol{w}^{\mathrm{S}}$ that satisfy the endosymbiont-level constraints in Eq. 1 as $\mathcal{A}(\boldsymbol{c}^{\mathrm{H}}; R^{\mathrm{S}})$.

$$\mathcal{A}(\boldsymbol{c}^{\mathrm{H}}; R^{\mathrm{S}}) := \left\{ \left(\boldsymbol{f}^{\mathrm{S}}, \boldsymbol{w}^{\mathrm{S}}\right) \;\middle|\; \begin{array}{l} \boldsymbol{0} = \boldsymbol{f}^{\mathrm{S}} + R^{\mathrm{S}}_{-m}\boldsymbol{w}^{\mathrm{S}} \\ \boldsymbol{f}^{\mathrm{S}} \leq \boldsymbol{c}^{\mathrm{H}} \\ \boldsymbol{w}^{\mathrm{S}} \geq \boldsymbol{0} \end{array} \right\}. \tag{S2}$$

Furthermore, we define the subset of $\mathcal{A}(\boldsymbol{c}^{\mathrm{H}}; R^{\mathrm{S}})$ with a specified growth rate $\mu$ as $\mathcal{A}_\mu$:

$$\mathcal{A}_\mu(\boldsymbol{c}^{\mathrm{H}}; R^{\mathrm{S}}) := \left\{ \left(\boldsymbol{f}^{\mathrm{S}}, \boldsymbol{w}^{\mathrm{S}}\right) \in \mathcal{A}(\boldsymbol{c}^{\mathrm{H}}; R^{\mathrm{S}}) \;\middle|\; R^{\mathrm{S}}_m \boldsymbol{w}^{\mathrm{S}} = \mu \right\}. \tag{S3}$$

Using the above definitions, the trilevel optimization we solved to obtain the phenotype of the host–endosymbiont pair is expressed as follows. The lowest layer of optimization is the endosymbiont growth maximization problem (Eq. 1), which defines the optimal growth rate $\mu^{\mathrm{S}*}$ given the amount of host-provided metabolites $\boldsymbol{c}^{\mathrm{H}}$ as:

$$\mu^{\mathrm{S}*}(\boldsymbol{c}^{\mathrm{H}}; R^{\mathrm{S}}) := \max_{(\boldsymbol{f}^{\mathrm{S}}, \boldsymbol{w}^{\mathrm{S}}) \in \mathcal{A}(\boldsymbol{c}^{\mathrm{H}}; R^{\mathrm{S}})} R^{\mathrm{S}}_m \boldsymbol{w}^{\mathrm{S}}. \tag{S4}$$

The middle layer of optimization corresponds to Eq. S1, which defines a unique $\boldsymbol{f}^{S*}$ given the optimal endosymbiont growth rate $\mu^{S*}(\boldsymbol{c}^H; R^S)$ as:

$$\left(\boldsymbol{f}^{S*}(\boldsymbol{c}^H; R^S), \boldsymbol{w}^{S*}(\boldsymbol{c}^H; R^S)\right) := \underset{(\boldsymbol{f}^S, \boldsymbol{w}^S) \in \mathcal{A}_{\mu^{S*}}(\boldsymbol{c}^H; R^S)}{\operatorname{argmin}} \|\boldsymbol{w}^S\|_2^2. \tag{S5}$$

Finally, the upper layer is the host growth maximization problem (Eqs. 2 and 3), which expresses both host adaptation through tuning $\boldsymbol{f}^H$, $\boldsymbol{w}^H$ and evolution through tuning $\boldsymbol{c}^H$. Note that these three layers do not coincide with the three levels in the main text: the lower and middle layers together correspond to level 1 (endosymbiont adaptation), and the upper layer combines levels 2 and 3 (host adaptation and evolution). Because host adaptation (level 2) and host evolution (level 3) in the main text maximize the same objective $\mu^H$, the two levels can be combined into a single maximization over $\boldsymbol{f}^H$, $\boldsymbol{w}^H$ and $\boldsymbol{c}^H$:

$$\begin{aligned} \max_{\boldsymbol{f}^H, \boldsymbol{w}^H, \boldsymbol{c}^H} \quad & \mu^H \\ \text{s.t.} \quad & \mu^H = R^H_m \boldsymbol{w}^H \\ & \boldsymbol{f}^{S*}(\boldsymbol{c}^H; R^S) = \boldsymbol{f}^H + R^H_{-m} \boldsymbol{w}^H \\ & \boldsymbol{f}^H \leq \boldsymbol{c}^E \\ & \boldsymbol{w}^H \geq \boldsymbol{0}. \end{aligned} \tag{S6}$$

#### S4 The Karush–Kuhn–Tucker (KKT) conditions

Host-level evolution makes the optimization problem complicated because $\boldsymbol{c}^H$, which affects the lower-level endosymbiont optimization problem, is a variable in the upper-level host optimization problem. Such nested problems are typically not solved directly; instead, the upper-level problem is optimized subject to the KKT conditions of the lower-level problems (Dempe, 2002). Similar to the Lagrange multiplier method, the KKT conditions give the conditions for optimality and can be used to convert an optimization problem into a set of algebraic constraints.

Specifically, the solution of the lower and middle layers of our optimization problem can be expressed as the intersection of two sets of solutions: the set of solutions that optimize endosymbiont growth (lower layer)

$$\mathcal{A}^*(\boldsymbol{c}^H; R^S) := \mathcal{A}_{\mu^{S*}(\boldsymbol{c}^H; R^S)}(\boldsymbol{c}^H; R^S) \tag{S7}$$

and the set of solutions that minimize the norm of the regulatory modes given any endosymbiont growth rate $\mu$, not necessarily the optimal one (middle layer)

$$\mathcal{B}(\boldsymbol{c}^H; R^S) := \bigcup_{\mu \in \mathbb{R}} \underset{(\boldsymbol{f}^S, \boldsymbol{w}^S) \in \mathcal{A}_\mu(\boldsymbol{c}^H; R^S)}{\operatorname{argmin}} \|\boldsymbol{w}^S\|_2^2 . \tag{S8}$$

Because the norm minimization is strictly convex, $\mathcal{A}^* \cap \mathcal{B} = \{(\boldsymbol{f}^{S*}, \boldsymbol{w}^{S*})\}$ is a single point, given $\boldsymbol{c}^H$ and $R^S$.

Thus, the upper-layer host optimization problem (Eq. S6) was solved subject to all KKT conditions of the lower- and middle-layer optimization problems, expressed below.

Any solution in $\mathcal{A}^*$ satisfies the following KKT conditions. We use the Lagrangian function $\mathcal{L}_{\mu^S}$ of the endosymbiont growth optimization problem max $\mu^S$ (Eq. 1) given as:

$$\mathcal{L}_{\mu^S}(\mu^S, \boldsymbol{w}^S) = \mu^S + \boldsymbol{\lambda}^1 \cdot \left( \begin{bmatrix} \boldsymbol{c}^H \\ 0 \end{bmatrix} + R^S \boldsymbol{w}^S - \begin{bmatrix} \boldsymbol{0} \\ \mu^S \end{bmatrix} \right) + \boldsymbol{\lambda}^2 \cdot \boldsymbol{w}^S, \tag{S9}$$

where $\boldsymbol{\lambda}^1$ and $\boldsymbol{\lambda}^2$ are the Lagrange multiplier vectors corresponding to the inequality constraints on nutrient uptake and the nonnegativity of the regulatory modes, respectively. Note that, using the mass conservation equations, we eliminated $\boldsymbol{f}^{\mathrm{S}}$ from Eq. S2 before obtaining the Lagrangian function. The KKT conditions are:

$$\begin{aligned}
&\text{Stationarity:}\\
&\quad 0 = \frac{\partial \mathcal{L}_{\mu^{\mathrm{S}}}}{\partial \mu^{\mathrm{S}}} = 1 - \lambda^1_m\\
&\quad \mathbf{0} = \frac{\partial \mathcal{L}_{\mu^{\mathrm{S}}}}{\partial \boldsymbol{w}^{\mathrm{S}}} = \boldsymbol{\lambda}^1 R^{\mathrm{S}} + \boldsymbol{\lambda}^2\\
&\text{Primal feasibility:}\\
&\quad \begin{bmatrix} \boldsymbol{c}^{\mathrm{H}} \\ 0 \end{bmatrix} + R^{\mathrm{S}}\boldsymbol{w}^{\mathrm{S}} - \begin{bmatrix} \mathbf{0} \\ \mu^{\mathrm{S}} \end{bmatrix} \geq \mathbf{0}, \quad \boldsymbol{w}^{\mathrm{S}} \geq \mathbf{0}\\
&\text{Dual feasibility:}\\
&\quad \boldsymbol{\lambda}^1 \geq \mathbf{0}, \quad \boldsymbol{\lambda}^2 \geq \mathbf{0}\\
&\text{Complementary slackness:}\\
&\quad \boldsymbol{\lambda}^1 \odot \left( \begin{bmatrix} \boldsymbol{c}^{\mathrm{H}} \\ 0 \end{bmatrix} + R^{\mathrm{S}}\boldsymbol{w}^{\mathrm{S}} - \begin{bmatrix} \mathbf{0} \\ \mu^{\mathrm{S}} \end{bmatrix} \right) = \mathbf{0}, \quad \boldsymbol{\lambda}^2 \odot \boldsymbol{w}^{\mathrm{S}} = \mathbf{0},
\end{aligned} \tag{S10}$$

where $\odot$ denotes the element-wise (Hadamard) product; $\boldsymbol{a} \odot \boldsymbol{b} := [a_1 b_1, a_2 b_2, \ldots, a_n b_n]^\top$. Roughly speaking, stationarity ensures optimality, primal feasibility ensures that the constraints are satisfied, dual feasibility fixes the direction of optimization, and complementary slackness handles whether each inequality is tight.

Any solution in $\mathcal{B}$ satisfies the following KKT conditions. We use the Lagrangian function $\mathcal{L}_{\boldsymbol{w}^{\mathrm{S}}}$ of the norm minimization problem given $\mu^{\mathrm{S}}$ (Eq. S5), which is:

$$\mathcal{L}_{\boldsymbol{w}^{\mathrm{S}}}(\boldsymbol{w}^{\mathrm{S}}; \mu^{\mathrm{S}}) = -\|\boldsymbol{w}^{\mathrm{S}}\|_2^2 + \boldsymbol{\lambda}^3 \cdot \left( \begin{bmatrix} \boldsymbol{c}^{\mathrm{H}} \\ 0 \end{bmatrix} + R^{\mathrm{S}}\boldsymbol{w}^{\mathrm{S}} - \begin{bmatrix} \mathbf{0} \\ \mu^{\mathrm{S}} \end{bmatrix} \right) + \boldsymbol{\lambda}^4 \cdot \boldsymbol{w}^{\mathrm{S}}, \tag{S11}$$

where $\boldsymbol{\lambda}^3$ and $\boldsymbol{\lambda}^4$ are Lagrange multiplier vectors, as in the previous case. Note the $-$ sign in the first term, because this is a minimization problem. Also, as we can again eliminate $\boldsymbol{f}^{\mathrm{S}}$ from Eq. S1 using the mass conservation equations, $\boldsymbol{w}^{\mathrm{S}}$ is the only variable in the Lagrangian function. Thus, the KKT conditions are:

$$\begin{aligned}
&\text{Stationarity:}\\
&\quad \mathbf{0} = \frac{\partial \mathcal{L}_{\boldsymbol{w}^{\mathrm{S}}}}{\partial \boldsymbol{w}^{\mathrm{S}}} = -2\boldsymbol{w}^{\mathrm{S}} + {R^{\mathrm{S}}}^\top \boldsymbol{\lambda}^3 + \boldsymbol{\lambda}^4,\\
&\text{Primal feasibility: (same as in the previous case)}\\
&\quad \begin{bmatrix} \boldsymbol{c}^{\mathrm{H}} \\ 0 \end{bmatrix} + R^{\mathrm{S}}\boldsymbol{w}^{\mathrm{S}} - \begin{bmatrix} \mathbf{0} \\ \mu^{\mathrm{S}} \end{bmatrix} \geq \mathbf{0}, \quad \boldsymbol{w}^{\mathrm{S}} \geq \mathbf{0}\\
&\text{Dual feasibility:}\\
&\quad \boldsymbol{\lambda}^3 \geq \mathbf{0}, \quad \boldsymbol{\lambda}^4 \geq \mathbf{0},\\
&\text{Complementary slackness:}\\
&\quad \boldsymbol{\lambda}^3 \odot \left( \begin{bmatrix} \boldsymbol{c}^{\mathrm{H}} \\ 0 \end{bmatrix} + R^{\mathrm{S}}\boldsymbol{w}^{\mathrm{S}} - \begin{bmatrix} \mathbf{0} \\ \mu^{\mathrm{S}} \end{bmatrix} \right) = \mathbf{0}, \quad \boldsymbol{\lambda}^4 \odot \boldsymbol{w}^{\mathrm{S}} = \mathbf{0}.
\end{aligned} \tag{S12}$$

#### S5 Numerical solution

##### S5.1 Levels 1 and 2

For a fixed $\boldsymbol{c}^{\mathrm{H}}$, the phenotype of a host–endosymbiont pair is obtained as a sequence of three convex programs: (i) $\max \mu^{\mathrm{S}}$ given $\boldsymbol{c}^{\mathrm{H}}$ (Eq. 1); (ii) $\min \|\boldsymbol{w}^{\mathrm{S}}\|_2^2$ given $\mu^{\mathrm{S}*}$ (Eq. S1); (iii) $\max \mu^{\mathrm{H}}$ given $\boldsymbol{f}^{\mathrm{S}*}$ (Eq. 2). The linear programs were solved with HiGHS 1.15.1 through its Python interface `highspy`, with the options `solver = ipm`, `run_crossover = off` and `presolve = off`.

The endosymbiont problem is homogeneous in $\boldsymbol{c}^{\mathrm{H}}$, so level 1 was solved on the normalized uptake limits $\boldsymbol{c}^{\mathrm{H}}/\|\boldsymbol{c}^{\mathrm{H}}\|_1$ and the solution rescaled afterwards. Step (ii) is strictly convex and was solved exactly with the dual active-set method as implemented in the `quadprog` package (v0.1.13) (Goldfarb and Idnani, 1983).

##### S5.2 Level 3

Using the KKT conditions above, the trilevel problem was written out as a single program that maximizes $\mu^{\mathrm{H}}$ subject to the KKT conditions of both max $\mu^{\mathrm{S}}$ and $\min\|\boldsymbol{w}^{\mathrm{S}}\|_2$. The complementary slackness constraints were expressed as special ordered sets of type 1 (SOS1), which allows them to be handled efficiently by branch and bound (Siddiqui and Gabriel, 2013). The resulting mixed-integer program was solved with CPLEX 22.2.0.0 through its Python API, with presolve disabled.

Branch and bound was stopped after $1.5\times10^5$ ticks (~10 min), set through the option `CPXPARAM_DetTimeLimit`. A truncated run returns a value that is feasible but not certified optimal, so the corresponding $\mu^{\mathrm{H}}$ is a lower bound on the true optimum. In the main design (5.95% of instances truncated), no instance with $r \leq 21$ was truncated, but some pairs reached the time limit at higher $r$ (Fig. S8). Such instances were more prevalent at higher $k$ and at higher $m$, where the programs are correspondingly harder.

In theory, the problem is always feasible, since $\boldsymbol{c}^{\mathrm{H}} = \boldsymbol{0}$ recovers the endosymbiont-free host ($\mu^{\mathrm{H}} = \mu_0^{\mathrm{H}}$). Indeed, the solver never reported infeasibility for the runs shown in the main figures, but did in a few supplementary runs, presumably for numerical reasons. Such instances were re-solved with numerical emphasis and, if needed, with presolve enabled. When the solver still reported infeasibility, the instance was discarded.

To make $\boldsymbol{c}^{\mathrm{H}}$ unique, we minimized the sum of the uptake limits $\|\boldsymbol{c}^{\mathrm{H}}\|_1$ in a second optimization subject to the same constraints and to $\mu^{\mathrm{H}} \geq \mu^{\mathrm{H}*} - \varepsilon$ ($\varepsilon = 10^{-9}$).

##### S5.3 Level 4: evolution of the number of regulatory modes

All populations shared a single randomly generated host ($m = 10$, $r^{\mathrm{H}} = 24$). Each population consisted of 25 host–endosymbiont lineages, each with its own endosymbiont $R^{\mathrm{S}}$ of $r_0 \in \{5, 15, 30, 45\}$ regulatory modes generated as in section S2. In each generation, each lineage mutated with probability 0.3, either gaining a new regulatory mode generated as in section S2 or losing a randomly chosen one with equal probability, so that mutation alone does not bias $r$. The next generation was formed by Wright–Fisher resampling of the lineages with probabilities proportional to $\mu^{\mathrm{H}}$ after host evolution (level 3). For each $r_0$, 160 independent populations were evolved.

#### S6 Quantities computed from the solutions

Classifying a metabolite as bounded or excreted, or a regulatory mode as active, means deciding whether a solver output is zero, so each classification needs a numerical tolerance; the tolerances used are given with the definitions.

A non-biomass metabolite $i$ is *bounded* when its uptake-limit constraint is active at the endosymbiont optimum, $c_i^{\mathrm{H}} - f_i^{\mathrm{S}*} \leq \tau$, and *unbounded* otherwise; $b$ and $u$ denote their counts, with $b + u = m - 1$. The tolerance combines an absolute and a relative part in the manner of `numpy.isclose`, $\tau = \tau_{\mathrm{abs}} + \tau_{\mathrm{rel}}\,|c_i^{\mathrm{H}}|$ with $\tau_{\mathrm{abs}} = 10^{-8}$ and $\tau_{\mathrm{rel}} = 10^{-6}$. After host evolution, components with $c_i^{\mathrm{H}} = 0$ occur; these are bounded when $f_i^{\mathrm{S}*} = 0$ and unbounded when $f_i^{\mathrm{S}*} < 0$, which is the classification consistent with the Jacobian decomposition below, since "bounded" is the active set of the constraint $f_i^{\mathrm{S}} \leq c_i^{\mathrm{H}}$.

A regulatory mode is *active* when $w_j^{\mathrm{S}*} > 10^{-8}$, and $r_+$ denotes their number; $R_{b+}^{\mathrm{S}}$ and $R_{u+}^{\mathrm{S}}$ are the corresponding $b \times r_+$ and $u \times r_+$ submatrices. The pseudoinverse $\left(R_{b+}^{\mathrm{S}}\right)^+$ was computed by singular value decomposition with singular values below $10^{-10}$ times the largest treated as zero. When $b = 0$ or $u = 0$ the sensitivity matrix $S$ is empty and $s = 0$. The expectations in the definitions of $\alpha$ and $s$ are taken over the sampled internal environments $\boldsymbol{c}^{\mathrm{H}}$ of a given $R^{\mathrm{S}}$.

The number of excreted metabolites is $\#\{i : f_i^{\mathrm{S}*} < -10^{-8}\}$, using the convention that $\boldsymbol{f}^{\mathrm{S}}$ is a net uptake, so that negative components are excretions into the host.

#### S7 Derivation of the adaptability and sensitivity of endosymbiont metabolism

This section gives in full the approximation summarized in *Materials and Methods*.

The nested optimization precludes direct calculation of how the endosymbiont's metabolic exchanges $\boldsymbol{f}^{\mathrm{S}}$ change with respect to the host-provided metabolites $\boldsymbol{c}^{\mathrm{H}}$. To make this calculation possible, we approximated $\boldsymbol{f}^{\mathrm{S}}$ as an explicit function of $\boldsymbol{c}^{\mathrm{H}}$. Since endosymbiont growth increases with the consumption of host-provided resources, endosymbiont growth maximization can be interpreted as maximizing the consumption of the available resources. We approximated this by minimizing the difference between the maximum rates at which the endosymbiont can take up each metabolite from the host ($\boldsymbol{c}^{\mathrm{H}}$) and its actual net uptake rates ($\boldsymbol{f}^{\mathrm{S}} = -R^{\mathrm{S}}_{-m}\boldsymbol{w}^{\mathrm{S}}$).

This approximation requires distinguishing two types of metabolite by their constraint status. Bounded metabolites reach their constraint limits in the optimization and are therefore completely utilized by the endosymbiont, while unbounded metabolites remain below their limits. The distinction is crucial because (i) endosymbiont metabolism responds to changes in the bounded components of $\boldsymbol{c}^{\mathrm{H}}$ (denoted $\boldsymbol{c}^{\mathrm{H}}_b$), whereas (ii) increasing the availability of an unbounded metabolite has no effect on the optimal endosymbiont metabolism, because being unbounded implies that the metabolite is already in excess.

Accordingly, the optimal metabolic exchanges are approximated as

$$\boldsymbol{f}^{\mathrm{S}*}(\boldsymbol{c}^{\mathrm{H}}; R^{\mathrm{S}}) \approx -R^{\mathrm{S}}_{-m} \underset{\boldsymbol{w}^{\mathrm{S}}}{\operatorname{argmin}} \left\|\boldsymbol{c}^{\mathrm{H}}_b + R^{\mathrm{S}}_b \boldsymbol{w}^{\mathrm{S}}\right\|_2 = R^{\mathrm{S}}_{-m}\left(R^{\mathrm{S}}_b\right)^+ \boldsymbol{c}^{\mathrm{H}}_b, \tag{S13}$$

where $b$ in the subscripts denotes the row subset corresponding to the bounded metabolites. For a nonsingular square matrix $A$, the solution of $\underset{\boldsymbol{x}}{\operatorname{argmin}}\|\boldsymbol{y} - A\boldsymbol{x}\|_2$ is $\boldsymbol{x} = A^{-1}\boldsymbol{y}$, at which the $\mathcal{L}^2$-norm is zero. For a general rectangular matrix $A$, $\boldsymbol{x} = A^{+}\boldsymbol{y}$ minimizes the residual and, among the minimizers, has the smallest norm, where the superscript + denotes the Moore–Penrose pseudoinverse. The minimum-norm property matches the tie-breaking rule used in the model itself (Eq. S1), so that under this approximation, inspired by the toolbox model (Tikhonov et al., 2020), the optimal weights of the endosymbiont's regulatory modes are $\boldsymbol{w}^{\mathrm{S}*} \approx -\left(R^{\mathrm{S}}_b\right)^+ \boldsymbol{c}^{\mathrm{H}}_b$.

Under small perturbations of the bounded metabolites $\boldsymbol{c}^{\mathrm{H}}_b$, regulatory modes that are not used by the endosymbiont ($w^{\mathrm{S}}_j = 0$) will typically remain unused, and only the active modes ($w^{\mathrm{S}}_j > 0$) respond and thereby influence $\boldsymbol{f}^{\mathrm{S}*}$. We therefore restrict attention to the column subset of $R^{\mathrm{S}}$ corresponding to the active regulatory modes, $R^{\mathrm{S}}_{+}$.

This formulation allows the partial derivative of $\boldsymbol{f}^{\mathrm{S}*}$ with respect to $\boldsymbol{c}^{\mathrm{H}}$ to be taken. With the metabolites reordered so that the first $b$ elements are the bounded non-biomass metabolites and the remaining $u$ are the unbounded ones, the Jacobian is approximated by

$$\begin{aligned} \frac{\partial \boldsymbol{f}^{\mathrm{S}*}}{\partial \boldsymbol{c}^{\mathrm{H}}_b} &\approx R^{\mathrm{S}}_{+}\left(R^{\mathrm{S}}_{b+}\right)^+ = \begin{bmatrix} R^{\mathrm{S}}_{b+} \\ R^{\mathrm{S}}_{u+} \end{bmatrix}\left(R^{\mathrm{S}}_{b+}\right)^+ = \begin{bmatrix} I_b \\ R^{\mathrm{S}}_{u+}\left(R^{\mathrm{S}}_{b+}\right)^+ \end{bmatrix}, \\ \frac{\partial \boldsymbol{f}^{\mathrm{S}*}}{\partial \boldsymbol{c}^{\mathrm{H}}_u} &= O_{m-1,u}, \end{aligned} \tag{S14}$$

where $u$ in the subscripts denotes the row subset corresponding to the unbounded metabolites (Fig. 5B). We used $I_b = R^{\mathrm{S}}_{b+}\left(R^{\mathrm{S}}_{b+}\right)^+$, that is, we assumed $R^{\mathrm{S}}_{b+}$ to have full row rank. This requires $r_+ \geq b$, which holds because a nondegenerate optimum of Eq. 1 has $r_+ = b$ and the minimum-norm selection (Eq. S1) can only raise $r_+$; the $b$ rows are then linearly independent except on a set of measure zero. The block $\partial\boldsymbol{f}^{\mathrm{S}*}/\partial\boldsymbol{c}^{\mathrm{H}}_u$ vanishes because the endosymbiont's metabolism is not affected by metabolites that are already in excess.

On this basis we decomposed the effect of host variation on endosymbiont metabolism, $\partial\boldsymbol{f}^{\mathrm{S}*}/\partial\boldsymbol{c}^{\mathrm{H}}_b$, into two components, the *adaptability* and the *sensitivity* of the endosymbiont:

$$\begin{aligned} \text{Adaptability } (\alpha) &:= \mathbb{E}[\sqrt{b}] & \left(\approx \mathbb{E}\left[\left\|\frac{\partial \boldsymbol{f}^{\mathrm{S}*}_b}{\partial \boldsymbol{c}^{\mathrm{H}}_b}\right\|_{\mathrm{F}}\right]\right) \\ \text{Sensitivity } (s) &:= \mathbb{E}\left[\|S\|_{\mathrm{F}}\right] & \left(\approx \mathbb{E}\left[\left\|\frac{\partial \boldsymbol{f}^{\mathrm{S}*}_u}{\partial \boldsymbol{c}^{\mathrm{H}}_b}\right\|_{\mathrm{F}}\right]\right) \end{aligned} \tag{S15}$$

Here $\mathbb{E}[\cdot]$ is the expectation over the random host-provided metabolite compositions $\boldsymbol{c}^{\mathrm{H}}$, $S := R^{\mathrm{S}}_{u+}\left(R^{\mathrm{S}}_{b+}\right)^+$ is the *sensitivity matrix*, and $\|S\|_{\mathrm{F}} := \sqrt{\sum_{i,j} S^2_{ij}}$ is its Frobenius norm. Note that $\|I_b\|_{\mathrm{F}} = \sqrt{b}$.

#### S8 Explanation for the quantitative relationship between the number of regulatory modes that gives peak host growth and the number of metabolites

Varying the number of non-biomass metabolites ($n$) and the number of regulatory modes ($r$) showed that the number of regulatory modes giving peak host growth increased linearly with the number of metabolites (Fig. 6C). This can be explained by the following argument, assuming that the tight constraints are randomly distributed between the two types of constraints (regulatory modes and metabolites).

The endosymbiont optimization (Eq. 1) has $r$ variables and $r + n$ inequality constraints: $r$ nonnegativity constraints on the regulatory modes and $n$ constraints on the uptake limits of the non-biomass metabolites. At a nondegenerate optimal vertex exactly $r$ constraints are tight (i.e., satisfied as equalities), each being either an inactive mode or a bounded metabolite, so $(r - r_+) + b = r$ and hence $r_+ = b$. If no constraint type is a priori more likely to be tight than the other, the tight set is a random $r$-subset of the $r + n$ constraints, giving $\mathbb{E}[b] = nr/(r + n)$ and $\mathbb{E}[u] = n^2/(r + n)$. Their product $n^3 r/(r + n)^2$ is maximized at $r = n$ for any $n$, which accounts for the linear relationship, and there half of the tight constraints come from each type, so that $r \approx 2b$.

This argument is approximate in three respects. First, it maximizes $\mathbb{E}[u]\,\mathbb{E}[b]$ rather than $\mathbb{E}[ub]$; the two differ by $\mathrm{Var}(b) = r^2 n^2/((r + n)^2(r + n - 1))$, which at $r = n$ is $O(n)$ against a product of $O(n^2)$, so the correction is of relative order $1/n$ and shifts the maximum slightly below $r = n$. Second, it assumes that every inequality in Eq. 1 is equally likely to be tight at the optimum, irrespective of whether it constrains a mode or a metabolite. This neglects the structure of $R^{\mathrm{S}}$, in particular that reaction directions are assigned randomly, so that metabolites differ in how readily they can be driven to their bounds. Third, the model does not use the vertex solution directly: the minimum-norm selection (Eq. S1) distributes weight among regulatory modes that are equally optimal, so the realized number of active modes satisfies $r_+ \geq b$ rather than $r_+ = b$. More modes are therefore required to reach a given $b$, which may contribute to the observed peak lying slightly above $r = n$.

A further caveat applies to the largest values of $m$. Trilevel problems that reached the time limit were assigned the best feasible solution found by the solver, which is a lower bound on the optimum (Fig. S8). Such cases became more frequent at larger $m$ and $k$; because they were concentrated at larger $r$, they would bias the estimated peak downward at large $m$.

#### S9 Statistical analysis

All statistical analyses were performed in R (v4.3.3). Unless stated otherwise, curves show the mean over host–endosymbiont pairs at each $r$. The $r$ at which a curve peaks ($r^*$) is the $r$ with the largest mean (ties broken toward the smaller $r$); its 95% confidence interval was obtained by a percentile bootstrap ($B = 2000$), resampling the endosymbionts ($R^{\mathrm{S}}$) with replacement independently within each $r$. Rank correlations are Spearman's $\rho$ (`cor.test`); partial correlations controlling for $r$ were computed as the correlation between the residuals of the ranks of both variables regressed on the rank of $r$. Linear relationships were fitted by ordinary least squares (`lm`), and logistic regressions with `glm`; where $r$ was controlled for, it entered as a natural cubic spline with 4 degrees of freedom. Highest density regions (Figs. 3C and 4C) were obtained from a Gaussian kernel density estimate with a fixed isotropic bandwidth ($h = 2 \times 10^{-3}$ and 0.2, respectively; standard deviation $h/4$) evaluated on a $256 \times 256$ grid, as the smallest density contour enclosing 25, 50 and 75% of the probability mass. Pairs for which the host-evolution problem was infeasible were excluded, and problems that reached the solver time limit were represented by the best solution found.

##### S9.1 Definition of the standard deviation of the endosymbiont metabolic exchanges

For each endosymbiont $R^{\mathrm{S}}$, the variation of its metabolic exchanges was computed across the $K$ (= 100) host-provided compositions $\boldsymbol{c}^{\mathrm{H}(k)}$ ($k = 1, \ldots, K$). Let $f_j^{(k)}$ denote the $j$-th component of $\boldsymbol{f}^{\mathrm{S}*}(\boldsymbol{c}^{\mathrm{H}(k)}; R^{\mathrm{S}})$, and let $\mathcal{K}_j^u$ be the set of compositions under which the $j$-th non-biomass metabolite is unbounded, classified as described in section S6. We define

$$\mathrm{SD}\left(\boldsymbol{f}_u^{\mathrm{S}*}\right) := \sqrt{\sum_{j=1}^{m-1} \frac{1}{\#\mathcal{K}_j^u} \sum_{k \in \mathcal{K}_j^u} \left(f_j^{(k)} - \bar{f}_j^u\right)^2}, \qquad \bar{f}_j^u := \frac{1}{\#\mathcal{K}_j^u} \sum_{k \in \mathcal{K}_j^u} f_j^{(k)}, \tag{S16}$$

where the term for metabolite $j$ is set to zero when $\#\mathcal{K}_j^u = 0$. That is, for each metabolite the variance is taken only over the compositions under which it is unbounded, and these variances are summed over metabolites before taking the square root. $\mathrm{SD}(\boldsymbol{f}_b^{\mathrm{S}*})$ is defined analogously, using the set $\mathcal{K}_j^b$ of compositions under which metabolite $j$ is bounded. This norm-type measure matches the Frobenius norms in the definitions of $\alpha$ and $s$.

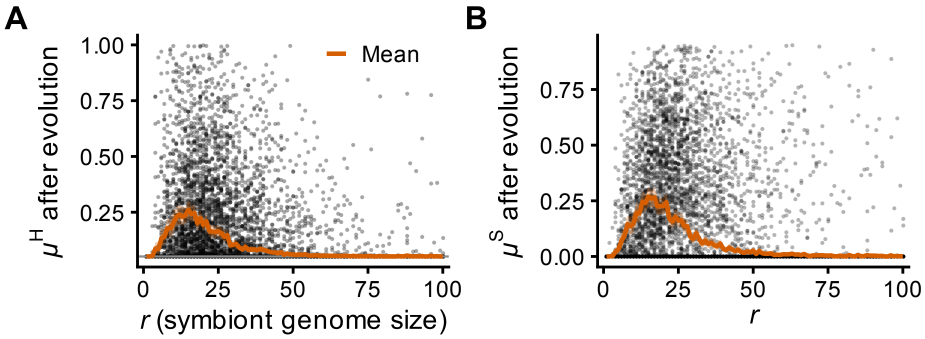


**Fig. S1. Evolution of mutualism with initially parasitic endosymbionts.** Random endosymbionts with varying genome sizes ($r$) were each assigned to a host with a fixed $R^{\rm H}$, and each host evolved to maximize its growth ($\mu^{\rm H}$) by tuning its internal environment ($\boldsymbol{c}^{\rm H}$) to exploit its endosymbiont. Each point corresponds to the growth ($\mu$) of a random host–endosymbiont pair with fixed $R^{\rm H}$ and random $R^{\rm S}$ as in Fig. 4AB, but only pairs with endosymbionts ($R^{\rm S}$) that were parasitic in the absence of host evolution (maximum $\mu^{\rm H}$ over the 100 random $\boldsymbol{c}^{\rm H}$ below the endosymbiont-free growth $\mu_0^{\rm H} = 0.052$; Fig. 3B) are shown (24,787 of 30,000 pairs). **(A)** $\mu^{\rm H}$ and **(B)** $\mu^{\rm S}$ after evolution. Orange lines show the mean for each $r$, with 95% bootstrap confidence intervals (shaded). The gray line in (A) shows the endosymbiont-free growth of the host.

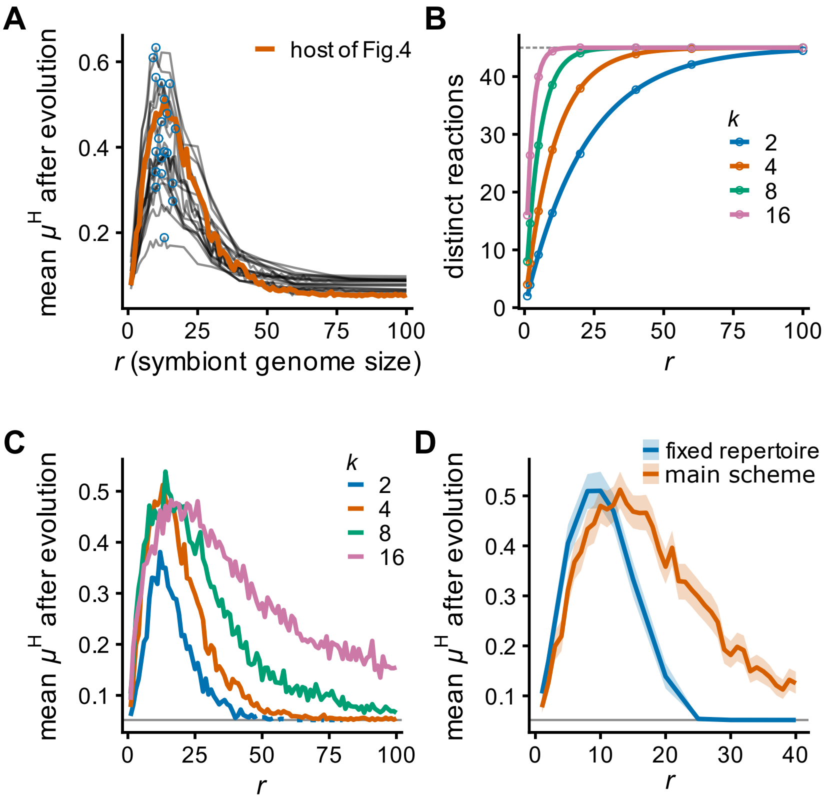


**Fig. S2. Robustness of the peak in $\mu^{\mathrm{H}}$ at intermediate $r$.** **(A)** Mean $\mu^{\mathrm{H}}$ after evolution against $r$ for 20 randomly generated hosts ($m = 10$, $k = 4$, $r^{\mathrm{H}} = 24$). The highlighted curve is the host of Fig. 4A. Hosts 0–9 were evolved with 300 random $R^{\mathrm{S}}$ at each $r = 1, \ldots, 100$, and hosts 10–19 with 100 random $R^{\mathrm{S}}$ at each of 15 values of $r$. Open circles mark the peak of each host (median $r^* = 12$, range 8–16). **(B)** Number of distinct reactions in a genome of $r$ regulatory modes with $k = 2, 4, 8$ or 16 reactions per mode, drawn from the ${}_{10}\mathrm{C}_2 = 45$ possible conversions. Lines show the theoretical expectation, and circles show direct simulations of the generation rule. The dashed line marks all 45 reactions. **(C)** Mean $\mu^{\mathrm{H}}$ after evolution against $r$ for $k = 2, 4$ (as in Fig. 4A), 8 and 16 (300 random $R^{\mathrm{S}}$ per $r$). **(D)** Mean $\mu^{\mathrm{H}}$ after evolution when a fixed set of 40 reactions is divided among the $r$ modes, so that the reaction repertoire does not change with $r$, compared with the scheme of Fig. 4A over the same range of $r$ (300 random $R^{\mathrm{S}}$ per $r$). The two curves peak at $r^* = 10$ (95% CI 8–10) and $r^* = 13$ (10–16), respectively. Shaded bands are 95% bootstrap confidence intervals. Gray lines in (C) and (D) show the endosymbiont-free growth of the host.

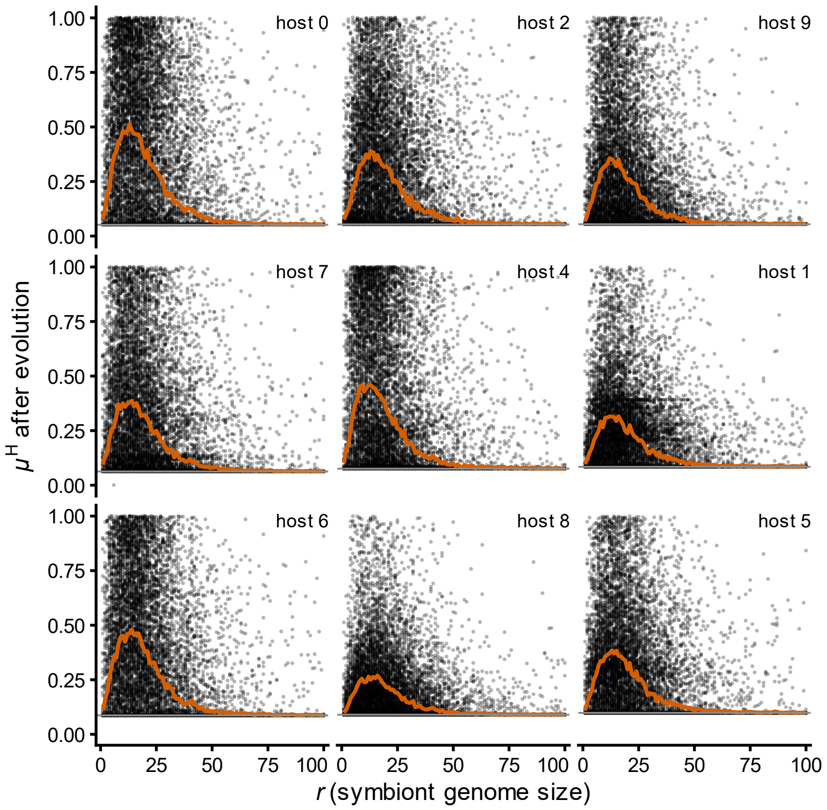


**Fig. S3. Variation in $\mu^{H}$ after evolution among host–endosymbiont pairs.** Each panel shows one representative host ($m$ = 10, $k$ = 4, $r^{H}$ = 24). Points are individual host–endosymbiont pairs (300 random $R^{S}$ at each $r = 1, \ldots, 100$), the orange line is the mean over the pairs at each $r$, and the gray line is the endosymbiont-free growth $\mu_0^{H}$ of that host (0.052–0.098), with panels ordered by increasing $\mu_0^{H}$. Host 0 is the host used in Fig. 4A. In every host, the mean $\mu^{H}$ peaked at an intermediate $r$ ($r^{*}$ = 12–16). One pair (host 7, $r$ = 6) returned $\mu^{H} = 0$ as an optimal solution, which is a numerical artifact of the MILP solver.

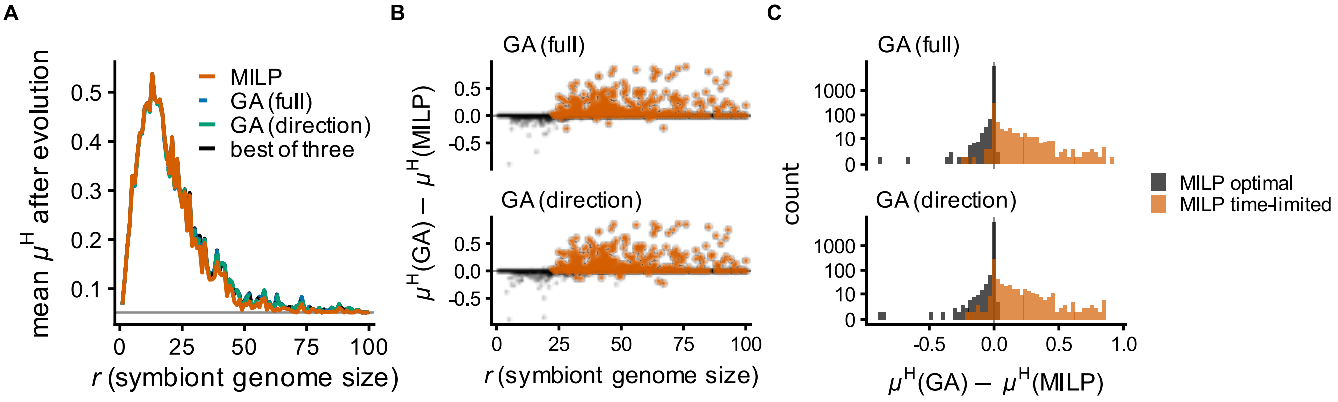


**Fig. S4. Robustness of the peak in $\mu^{\mathrm{H}}$ to the optimization method.** To test whether the peak in $\mu^{\mathrm{H}}$ at intermediate $r$ was an artifact of the MILP solver, we compared its results with those of two genetic algorithms (GAs) that do not rely on the MILP solver. Host evolution was solved again with a GA for the host and endosymbionts of Fig. 4A (100 random $R^{\mathrm{S}}$ per $r = 1, \ldots, 100$), and the results were compared with the MILP solutions pair by pair. GA (full) evolves $\boldsymbol{c}^{\mathrm{H}}$ directly (population 100, 300 generations); GA (direction) evolves only the direction of $\boldsymbol{c}^{\mathrm{H}}$ and line-searches its magnitude (population 100, 60 generations). The MILP reached its time limit in 574 of the 10,000 pairs (5.7%). **(A)** Mean $\mu^{\mathrm{H}}$ after evolution against $r$ for each method and for the best of the three methods in each pair. All four curves peak at $r^* = 13$ (95% bootstrap CI 10–16; 11–16 for GA (direction)). The gray line shows the endosymbiont-free growth of the host. **(B)** Difference in $\mu^{\mathrm{H}}$ between the GA and the MILP for each pair, against $r$. Gray points are pairs for which the MILP found the optimal solution (MILP optimal); orange points are pairs in which it reached its time limit (MILP time-limited). **(C)** Distributions of the differences in (B). Where the MILP was optimal, the differences are concentrated at zero. Where it reached its time limit, the GA found a higher $\mu^{\mathrm{H}}$ in 69.5% (GA (full)) and 60.5% (GA (direction)) of the pairs, with a mean difference of +0.106 and +0.105, respectively. GA (full) reached a host growth rate no more than $10^{-2}$ below that of the MILP in 98% of the 10,000 pairs.

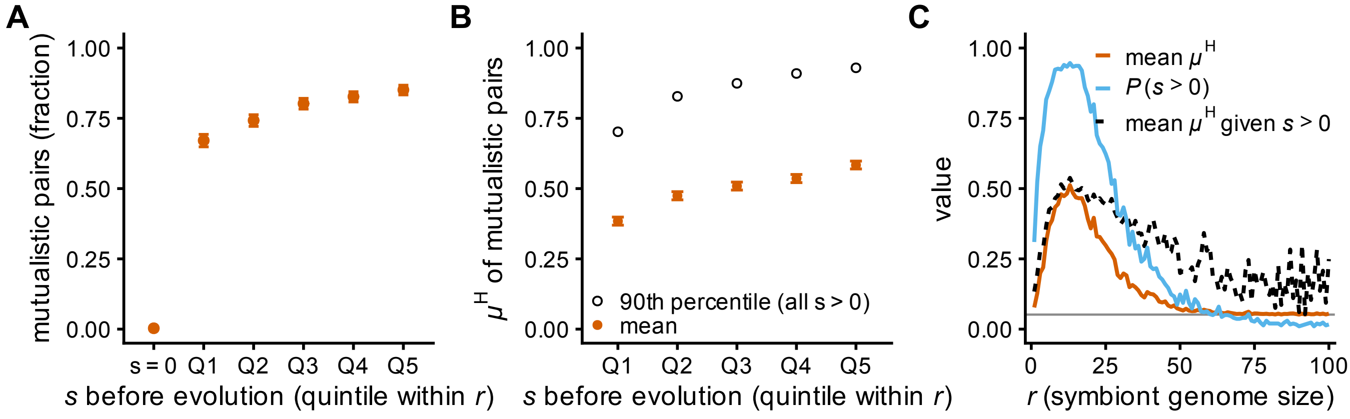


**Fig. S5. Sensitivity is required for mutualism but only modestly predicts its extent.** Host–endosymbiont pairs of Fig. 4A ($N$ = 30,000; 300 random $R^{\mathrm{S}}$ per $r = 1, \ldots, 100$). A pair was regarded as mutualistic when $\mu^{\mathrm{H}} > \mu_0^{\mathrm{H}} + 10^{-2}$ and $\mu^{\mathrm{S}} > 10^{-2}$ after host evolution. $s$ is the mean sensitivity of the endosymbiont over 100 random $\boldsymbol{c}^{\mathrm{H}}$ before host evolution, so $s > 0$ means that $s$ was positive for at least one $\boldsymbol{c}^{\mathrm{H}}$. **(A)** Fraction of mutualistic pairs among endosymbionts with $s = 0$ (21,300 pairs) and among endosymbionts with $s > 0$, divided into quintiles of $s$ within each $r$ (Q1–Q5, 1,700–1,775 pairs each). Error bars are exact binomial 95% confidence intervals. Only 0.3% of the pairs with $s = 0$ became mutualistic, compared with 68–88% of the pairs with $s > 0$; 6,935 of the 7,003 mutualistic pairs (99%) had $s > 0$. **(B)** Mean $\mu^{\mathrm{H}}$ after evolution of the mutualistic pairs in each quintile of (A) (filled circles, with 95% confidence intervals), and the 90th percentile of $\mu^{\mathrm{H}}$ after evolution among all pairs with $s > 0$ in the quintile (open circles). **(C)** Mean $\mu^{\mathrm{H}}$ after evolution (as in Fig. 4A), the fraction of endosymbionts with $s > 0$, and the mean $\mu^{\mathrm{H}}$ after evolution of the endosymbionts with $s > 0$, against $r$. All three peak at $r = 13$. The gray line shows the endosymbiont-free growth of the host.

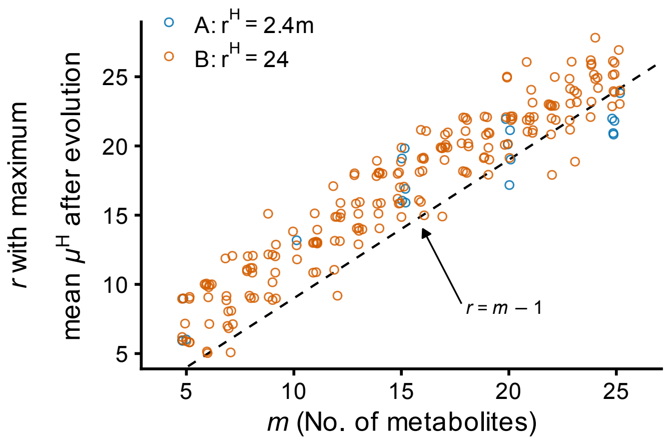


**Fig. S6. Optimal endosymbiont genome size when the host genome size scales with the number of metabolites.** As in Fig. 6C, each point shows, for one randomly generated host, the $r$ that achieved the maximum mean $\mu^{\mathrm{H}}$ after evolution, against the number of metabolites $m$. Orange circles are the hosts of Fig. 6C, with a fixed host genome size $r^{\mathrm{H}} = 24$ (10 hosts per $m = 5, 6, \ldots, 25$). Blue circles are hosts whose genome size scales with $m$, $r^{\mathrm{H}} = \mathrm{round}(2.4\,m)$ (5 hosts at each of $m = 5$, 10, 15, 20 and 25); at $m = 10$ the two designs coincide ($r^{\mathrm{H}} = 24$). Hosts were evolved over $r$ as in Fig. 6C. The dashed line shows $r = m - 1$. Points are jittered by at most 0.2 on both axes.

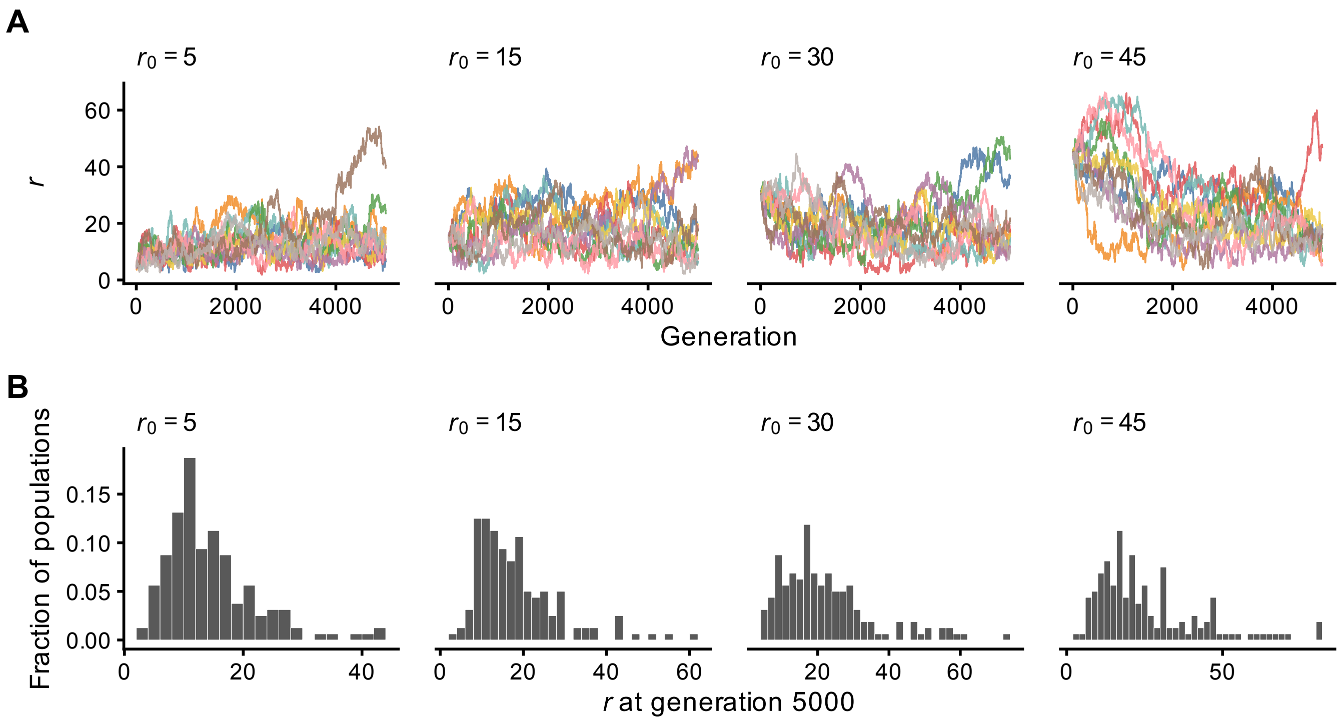


**Fig. S7. Evolution of the number of regulatory modes $r$ in replicate populations.** For each initial value $r_0 \in \{5, 15, 30, 45\}$, 160 independent Wright–Fisher populations (25 lineages each) were evolved, and $r$ was recorded every generation until generation 5000. **(A)** $r$ against generation for 10 randomly chosen populations per $r_0$. Colors distinguish the populations within each panel. **(B)** Distribution of $r$ at the most recent generation recorded for each population (160 populations per $r_0$), shown as the fraction of populations within each $r_0$.

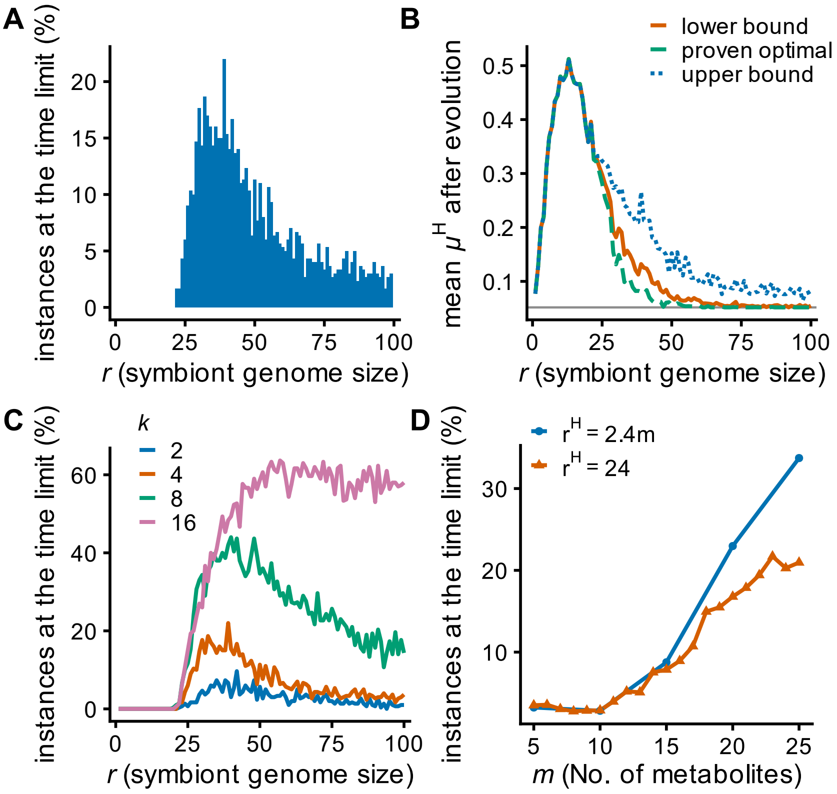


**Fig. S8. Effect of the solver time limit on host evolution.** **(A)** Percentage of host-evolution problems that reached the deterministic time limit of the MILP solver ($1.5 \times 10^5$ ticks; section S5.2), against $r$ ($m = 10$, $k = 4$, $r^H = 24$, 300 random $R^S$ per $r$, as in Fig. 4A). Overall, 6.0% of the problems (1,785 of 30,000) reached the limit. None did at $r \leq 21$, and the percentage was highest at $r = 39$ (22.0%). **(B)** Results using the best feasible (not certified optimal) solutions from the solver to estimate $\mu^H$ after evolution. Mean $\mu^H$ after evolution against $r$ under three treatments of time-limited problems: the best feasible solution found by the solver (lower bound; used in all other figures); that solution multiplied by $1 + g$, where $g$ is the relative MIP gap reported by the solver, capped at the model's maximum $\mu^H = 1$ (upper bound); and only the problems solved to optimality. All three curves peak at $r^* = 13$, and the upper and lower bounds differ by at most 0.13 (at $r = 39$). The gray line shows the endosymbiont-free growth of the host. **(C)** Percentage of problems that reached the time limit against $r$, for endosymbionts generated with $k = 2, 4, 8$, or 16 reactions per regulatory mode (300 random $R^S$ per $r$; same simulations as Fig. S2C). Overall, 2.3%, 6.0%, 20.8%, and 40.8% of the problems reached the limit, respectively. **(D)** Percentage of problems that reached the time limit against the number of metabolites $m$, for hosts with $r^H = 24$ (10 hosts per $m$, as in Fig. 6C) and for hosts with $r^H = \mathrm{round}(2.4\,m)$ (5 hosts at each of $m = 5, 10, 15, 20, 25$). Because the values of $r$ sampled differ between $m$, the percentages were computed only over the values sampled for every host and $m$ ($r \in \{1, 2, 3, 5, 8, 10, 12, 15, 20, 25, 30, 40, 60, 100\}$).

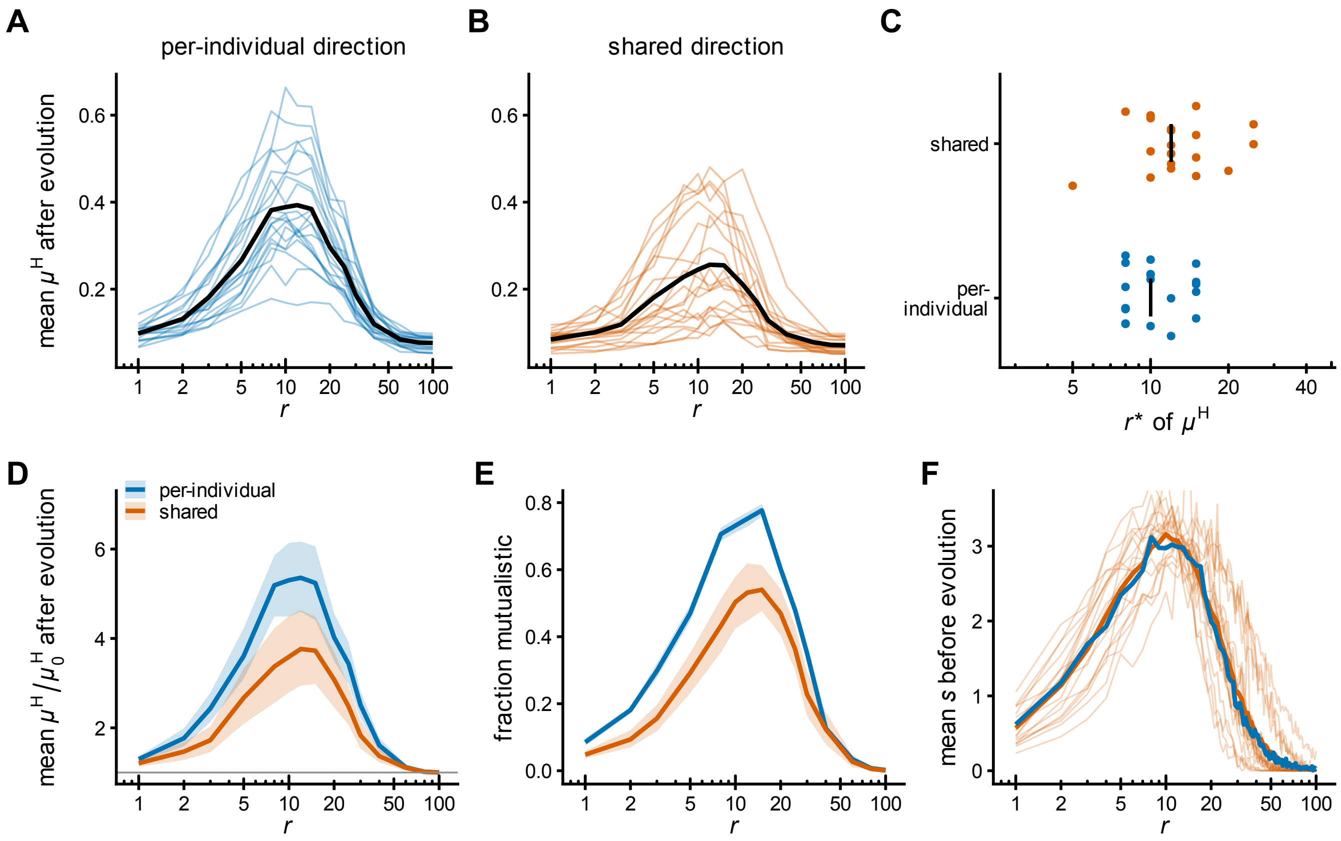


**Fig. S9. Host growth after evolution when reaction directions are shared between host and endosymbionts.** In the per-individual scheme (main text), the direction of each reaction is drawn independently for every host and every endosymbiont. In the shared scheme, one direction per metabolite pair is drawn per world and used by the host and all endosymbionts of that world. $m = 10$, $k = 4$, $r^H = 24$. For the shared scheme, 20 worlds were generated, each with one host. Twenty hosts for each scheme were evolved with 100 endosymbionts at each of 15 values of $r$. The $r$ axis is logarithmic. **(A, B)** Mean $\mu^H$ after evolution against $r$ for the per-individual scheme (A) and the shared scheme (B). Thin lines, individual hosts (A) or worlds (B); black line, mean over hosts or worlds. **(C)** Genome size $r^*$ maximizing the mean $\mu^H$, per host (blue) and per world (orange), jittered vertically. Black bars, medians. **(D)** Mean $\mu^H/\mu_0^H$ after evolution, where $\mu_0^H$ is the host growth rate without an endosymbiont. Gray line, $\mu^H/\mu_0^H = 1$. **(E)** Fraction of evolved pairs that are mutualistic ($\mu^H > \mu_0^H + 10^{-2}$ and $\mu^S > 10^{-2}$). In D and E, lines are means over hosts or worlds, and bands are bootstrap 95% confidence intervals with hosts or worlds resampled. **(F)** Mean sensitivity $s$ before evolution, averaged over $\boldsymbol{c}^H$ per endosymbiont and then over endosymbionts, at $r = 1, \ldots, 100$. Thin orange lines, individual worlds (100 endosymbionts $\times$ 10 $\boldsymbol{c}^H$ per $r$); thick orange line, mean over worlds; blue line, per-individual scheme (300 endosymbionts $\times$ 100 $\boldsymbol{c}^H$ per $r$, as in Fig. 6B). The $y$-axis is truncated at 1.05 times the 99th percentile of the plotted values.

**Table S1.** Symbols in this study

| Symbol | Description | Type or default value |
|---|---|---|
| **Model** | | |
| $\mu^{X}$ | Growth or biomass production rate of $X$ (H: host or S: endosymbiont) | optimized by X |
| $\mu_0^{H}$ | Endosymbiont-free growth of host | $\mu_0^{H} \in [0.05, 0.1]$ |
| $R^{X}$ | Regulatory matrix of $X$ | randomly generated as genotype |
| $\boldsymbol{w}^{X}$ | Weights of the regulatory modes (columns of $R^{X}$) | optimized by $X$ |
| $\boldsymbol{c}^{E}$ | Upper bound on the rate at which hosts can take up each metabolite from the environment | fixed |
| $\boldsymbol{c}^{H}$ | Upper bound on the rate at which endosymbionts can take up each metabolite from the host | optimized by host |
| $\boldsymbol{f}^{H}$ | Net uptake rate of metabolites from the environment by host | optimized by host |
| $\boldsymbol{f}^{S}$ | Net uptake rate of metabolites by endosymbiont | optimized by endosymbiont |
| $r$ | Number of independent regulatory modes of endosymbiont | varied from 1 to 100 |
| $m$ | Number of metabolites | 10 (unless stated otherwise) |
| $n$ | Number of non-biomass metabolites | $m - 1$ |
| $k$ | Number of reactions per regulatory mode | 4 (unless stated otherwise) |
| $r^{H}$ | Number of regulatory modes of host | 24 (unless stated otherwise) |
| **Analysis** | | |
| $b$ | Number of bounded metabolites | |
| $u$ | Number of unbounded non-biomass metabolites | |
| $R_b^{X}$ | Row subset of $R^{X}$ with only rows of bounded metabolites | |
| $R_u^{X}$ | Row subset of $R^{X}$ with only rows of unbounded metabolites | |
| $X^{+}$ | Pseudoinverse of $X$ | |
| $R_+^{X}$ | Column subset of $R^{X}$ with only columns of active regulatory modes | |
| $r_+$ | Number of active regulatory modes | |
| $r^*$ | Number of regulatory modes that maximizes the mean $\mu^{H}$ after host evolution | |
| $\alpha$ | Adaptability of endosymbiont metabolism to changes in host-provided metabolites | |
| $S$ | Sensitivity matrix of endosymbiont metabolism | |
| $s$ | Sensitivity (scalar), $\mathbb{E}\left[\|S\|_F\right]$ | |

### SI References


Cantalapiedra, C. P., Hernández-Plaza, A., Letunic, I., Bork, P., and Huerta-Cepas, J. eggNOG-mapper v2: Functional Annotation, Orthology Assignments, and Domain Prediction at the Metagenomic Scale. *Molecular Biology and Evolution*, 38(12):5825–5829, 2021. doi: 10.1093/molbev/msab293.

Dempe, S. *Foundations of Bilevel Programming*, volume 61. Kluwer Academic Publishers, Boston, 2002. ISBN 978-1-4020-0631-9. doi: 10.1007/b101970.

Forni, G., Martelossi, J., Morel, B., Pistone, D., Bandi, C., and Montagna, M. Large-scale phylogenomics reveals convergent genome evolution across repeated transitions to endosymbiosis in Enterobacterales. *Molecular Phylogenetics and Evolution*, 217:108532, 2026. doi: 10.1016/j.ympev.2026.108532.

Goldfarb, D. and Idnani, A. A numerically stable dual method for solving strictly convex quadratic programs. *Mathematical Programming*, 27(1):1–33, 1983. doi: 10.1007/BF02591962.

Larsson, J., Nylander, J. A., and Bergman, B. Genome fluctuations in cyanobacteria reflect evolutionary, developmental and adaptive traits. *BMC Evolutionary Biology*, 11(1):187, 2011. doi: 10.1186/1471-2148-11-187.

Lhee, D., Ha, J.-S., Kim, S., Park, M. G., Bhattacharya, D., and Yoon, H. S. Evolutionary dynamics of the chromatophore genome in three photosynthetic *Paulinella* species. *Scientific Reports*, 9(1):2560, 2019. doi: 10.1038/s41598-019-38621-8.

Lomsadze, A., Gemayel, K., Tang, S., and Borodovsky, M. Modeling leaderless transcription and atypical genes results in more accurate gene prediction in prokaryotes. *Genome Research*, 28(7):1079–1089, 2018. doi: 10.1101/gr.230615.117.

Manzano-Marín, A. and Latorre, A. Snapshots of a shrinking partner: Genome reduction in *Serratia symbiotica*. *Sci. Rep.*, 6(1):32590, 2016. doi: 10.1038/srep32590.

Siddiqui, S. and Gabriel, S. A. An SOS1-based approach for solving MPECs with a natural gas market application. *Networks and Spatial Economics*, 13(2):205–227, 2013. doi: 10.1007/s11067-012-9178-y.

Siozios, S., Nadal-Jimenez, P., Azagi, T., Sprong, H., Frost, C. L., Parratt, S. R., Taylor, G., Brettell, L., Liew, K. C., Croft, L., King, K. C., Brockhurst, M. A., Hypša, V., Novakova, E., Darby, A. C., and Hurst, G. D. D. Genome dynamics across the evolutionary transition to endosymbiosis. *Current Biology*, 34(24):5659–5670.e7, 2024. doi: 10.1016/j.cub.2024.10.044.

Tikhonov, M., Kachru, S., and Fisher, D. S. A model for the interplay between plastic tradeoffs and evolution in changing environments. *Proc. Natl. Acad. Sci. U. S. A.*, 117(16):8934–8940, 2020. doi: 10.1073/pnas.1915537117.
